# Reuse of learned subtask representations during human planning

**DOI:** 10.64898/2026.10.12.758531

**Authors:** Leo Chi U Seak, Lennart Luettgau, Maria K. Eckstein, Rani Moran, Raymond J. Dolan

## Abstract

The notion of a compositional reuse of previously learned problem solutions predicts that the human brain should reinstate such learned subtask representations during planning. Here, we test this in a value-ranking task implemented in an online and a magnetoencephalography (MEG) experiment, respectively. Participants first learned pairwise rankings and subsequently solved problems where these learned rankings could either be reused or required reordering. Behaviourally, in the online experiment, we show the availability of a reusable subtask structure improved accuracy, with computational modelling indicating a greater weighting of subtask-level value when participants could avail of this learnt structure. In MEG, during planning, we show enhanced reactivation of task-relevant subtask representations in the reusable condition, and particularly so early in the experiment. In a subsequent probe task, we show subtask-identity representations generalised to a novel context and that stimuli belonging to the same learned subtask manifest an increase in their neural similarity. Temporally delayed linear modelling (TDLM) revealed forward and backward sequential reactivation at a subtask level, alongside backward stimulus-level replay, with subtask sequenceness varying according to task context. The findings provide convergent behavioural, computational and neural evidence for reactivation and reuse of learnt relational structures for novel problem solving.

## Introduction

Humans show exceptional flexibility in performing complicated tasks across changing needs and demands (Tenenbaum et al., 2011; Reverberi et al., 2012; Ito et al., 2022; Schwartenbeck et al., 2023). Historically, humans variously composited stones for tools (Stout, 2011), sounds for languages (Botha and Knight, 2009), and plant material for fire (Davis et al., 2026).

Likewise, in daily life, tasks as mundane as cooking a simple dish invariably involve preparing, previously learned, individual elements separately and combining these to complete a meal, as well as availing of these separate elements when creating entirely new dishes in the future. This ability to flexibly reuse elements of acquired knowledge is among the defining features of human intelligence, and remains an aspiration in the design of agents endowed with artificial general intelligence (AGI) (Legg and Hutter, 2007; Sarikaya, 2025).

Compositionality involves the construction of novel representations from a finite set of familiar elements (Chollet, 2019; Wyble et al., 2025), where simpler elements provide core building blocks. Compositional abilities are not a sole prerogative of humans, other species, for example rhesus monkeys, flexibly compose tasks from subtask elements (Tafazoli et al., 2026). Nevertheless, how these subtask representations are learned, and made available for use during planning of multifaceted complex tasks remains unknown.

Two candidate neural mechanisms can be considered as supporting subtask utilisation. Prior human and non-human primate data spanning visual, language and navigation tasks suggest context-independent reactivation of ‘schema’ neurons that share a single conceptual feature (Baraduc et al., 2019; Bernardi et al., 2020; Dijksterhuis et al., 2024; Farzanfar et al., 2023; Mackay et al., 2024; Quiroga et al., 2005). In essence this involves reactivation of abstract, context-independent, subtask representations. Indeed, non-invasive human neuroimaging data show that distinct stimuli that share conceptual features express a common neural pattern (Favila et al., 2020; Lee et al., 2026; Luettgau et al., 2025). Whether learned subtask solutions are selectively reinstated during planning remains less clear.

A second, not unrelated, mechanism invokes rapid sequential reactivation of learned representations, as widely implicated in learning and planning (Buhry et al., 2011; Diba and Buzsáki, 2007; Foster and Wilson, 2006; Mattar and Daw, 2018). Relevant here are human magnetoencephalography (MEG) studies reporting replay of factorised task-relevant features, such as stimulus position and identity (Liu et al., 2019, 2021b; Schwartenbeck et al., 2023; Wimmer et al., 2023). Whether replay operates over learned subtask solutions, rather than over stimulus elements alone, remains an open question.

To examine subtask learning and reutilisation, we embedded a value ranking task within an experimental context where learned subtask elements could, or could not, be repurposed. We hypothesised that compositional planning, involving reuse of learned subtask components, would manifest as enhanced behavioural accuracy compared to a situation where subtask representations hold no such benefit. Likewise, at a neural level we hypothesised enhanced reactivation, and replay, of subtask representations within a context where these facilitate task solution. We report enhanced behavioural performance when subtask solutions could be repurposed to solve a current task, an effect captured within a reinforcement learning model involving a weighted mixture of subtask-level and stimulus-level value. In keeping with this, enhanced reactivation of subtask representations was evident in the very same context where these enabled a task solution, and when these enabled a solution to a novel out-of-sample problem.

## Results

### Task design

In a novel value ranking task (Fig. 1A) participants first learned a value ranking (high-to-low) of two stimuli (i.e. a subtask) through direct feedback, with values randomly assigned to stimuli on every trial (from −10 to +10), for each of three distinct subtasks. Thus, when two stimuli appeared on screen, participants were instructed to select a high-to-low ranking order (by pressing buttons), after which the value of each stimulus was revealed with subjects informed if the ranking was correct (Fig. 1A, bottom left). Two consecutive correct rankings were required before advancing to learning the next subtask. After learning three distinct subtasks, participants were then required to solve a 4-stimuli (from two randomly selected subtasks) ranking problem, which could follow a high-to-low or low-to-high value. In this planning phase, the four constitutive stimuli and ranking instruction were displayed.

**Figure 1.**
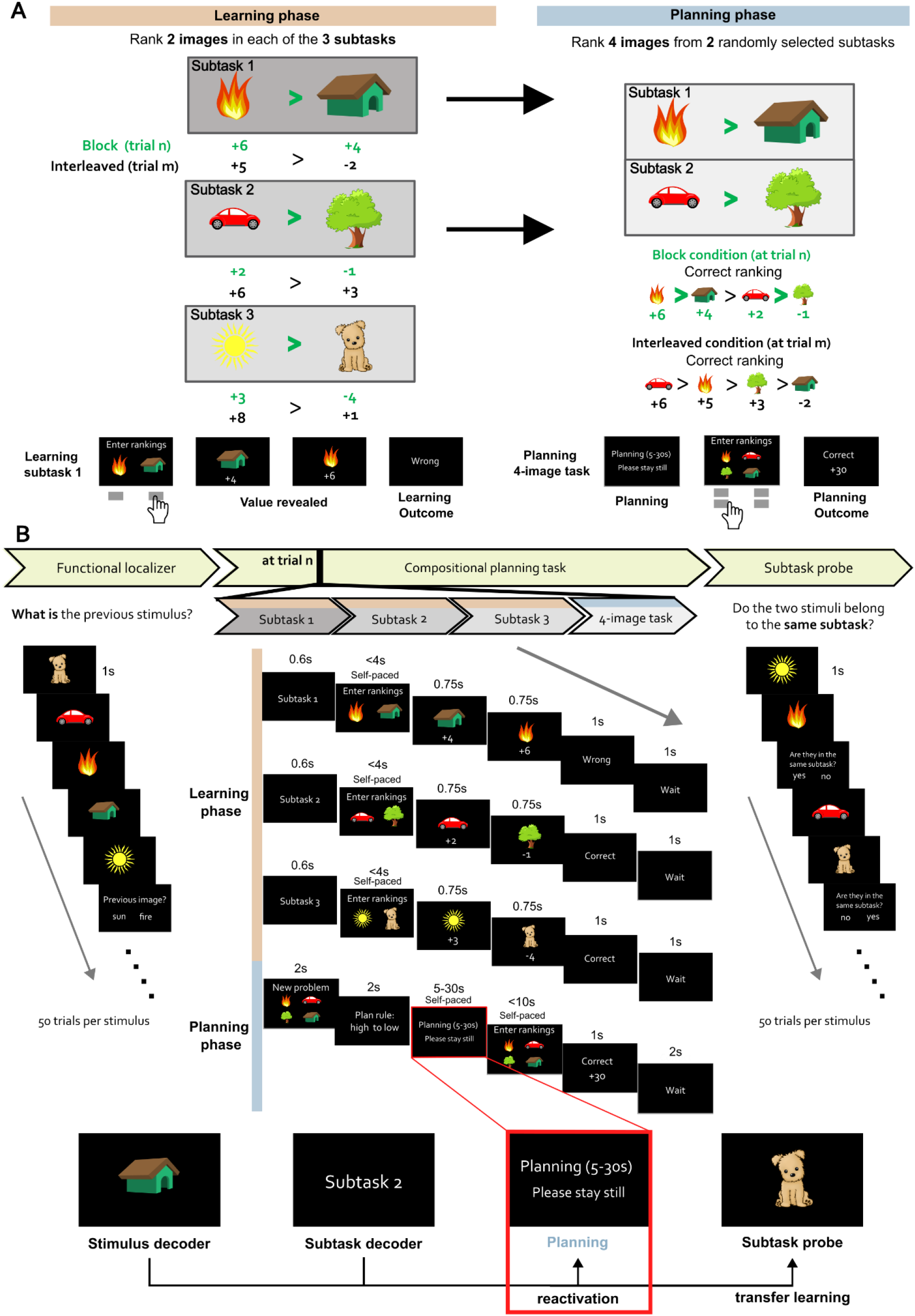
Task design. (A) Problem-solving task. Participants were tasked to rank order visual stimuli according to an assigned value (from −10 to +10), learning rankings of two discrete stimuli within each of three designated subtasks. Subsequently, they were presented with a 4-stimuli ranking problem (planning task based on two subtasks randomly selected at each trial). Throughout the experiment, subtasks contained the same two stimuli (but differed across participants) whose values changed randomly at every trial. Unbeknown to participants, there were two distinct task conditions. A block condition (trial n in the example) wherein participants could reuse both value-image associations and ranking information (subtask information); and an interleaved condition (trial m in the example), where they could only avail of individual value-image associations. Specifically, despite fire>house and car>tree in value for both trial n and trial m, the former ranking can be reused directly to solve the task in the block condition (trial n), whereas in the interleaved condition (trial m) such ranking information cannot be reused. Bottom panel shows an on-screen sequence for learning subtask 1 (left) and a planning 4-stimuli ranking problem (right). In a learning phase, the two stimuli are presented, participants select them in high-to-low ranked order, with the value of each stimulus followed by ranking feedback (correct/incorrect). To advance to the next subtask required two consecutive correct rankings. In the planning phase, the four stimuli and the ranking rule are shown, after which participants plan and respond by selecting the images in the required order. (B) MEG scanning protocol. MEG data were acquired throughout the experiment, which comprised a stimulus functional localiser; a problem-solving task; and a post-experiment subtask probe. The localiser task involved presenting one of the six visual images to participants in pseudorandom order. In the ranking task itself, participants learned three subtask rankings. To proceed to the next subtask, they needed to show subtask knowledge by answering a subtask ranking correctly (e.g. fire>house in the subtask 1 of the figure) twice in a row. A 30-pence bonus (10 pence for the online experiment) was provided if the 4-stimuli ranking problem was answered correctly. As shown in the bottom of the panel, primary time-points of interest for MEG data analyses related to (i) the time when participants planned a solution to a 4-image ranking problem (5-30s; red rectangle in the figure) and (ii) the subtask probe session when participants were shown two consecutive pseudorandom stimuli and tasked to indicate whether or not they belonged to a unique subtask. The stimulus and subtask decoders were respectively trained at the functional localiser (when stimulus images were shown) and the subtask learning phase (when the subtask text was shown).

Participants provided a full ranking by selecting images in the required order (Fig. 1A, bottom right). A bonus monetary reward (30 pence for the MEG experiment; 10 pence for the online experiment) was provided for a correct answer to this 4-stimuli problem.

Unbeknown to participants, we imposed two distinct task conditions, comprising a block and interleaved condition respectively. In the block condition, learned subtask rankings could be concatenated to provide a 4-stimuli solution, in effect reusing previously learned subtasks. In the interleaved condition, only a single value-image association could guide choice. The online behavioural and MEG experiments were identical in structure, apart from MEG including a pre-experiment functional localiser task (at session beginning) and a post-experiment subtask probe (Fig. 1B), serving to train stimulus decoders and test whether subtask representations transfer to a new context respectively.

### Behavioural and modelling of compositional subtask reuse

In an online experiment (N=241), we tested for a behavioural signature of compositional planning involving subtask reuse. A second, follow-up experiment involved an in-lab MEG study (N=25) that used the exact same paradigm. Our overarching hypothesis was that in a block condition (see Figure 1A) there would be reuse of learned subtask solutions leading to higher behavioural accuracy and, in turn, this reuse would be linked to a greater neural subtask reactivation when this facilitated a problem solution.

In both the online and MEG experiments there were behavioural features consistent with task engagement including improved accuracy across trials (Fig. S1A). In the online experiment (Fig. S1B) there was an association between longer planning times and greater task accuracy, although planning times per se did not differ between block and interleaved conditions. Moreover, participants performed better when asked to rank from a high-to-low than from low-to-high ranking in the online experiment (Fig. S2), aligning with the native direction of the learned subtask rankings.

Consistent with our behavioural hypothesis, in the online experiment participants showed higher accuracy in the block compared to the interleaved condition (43.5% vs 36.4%, z = 5.493, p <0.001; Wilcoxon signed-rank test, Fig. 2A) with a marginally significant effect evident for the MEG experiment (39.6% vs 33.6%, z = 1.943, p = 0.0520, Fig. 2A inset). Both exceeded a chance level of 0.042 (1/24 possible orders).

**Figure 2.**
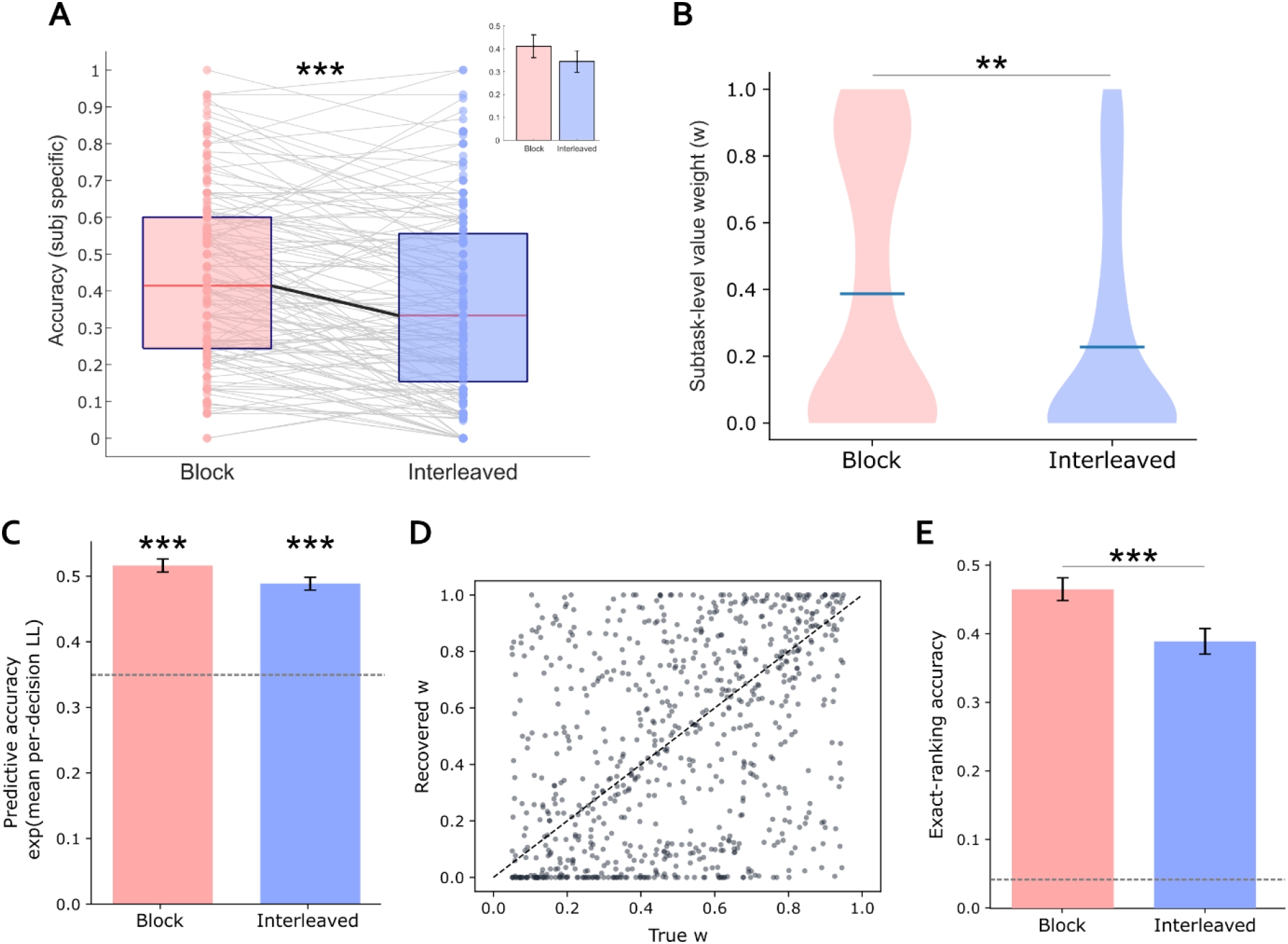
Behaviour and models. We fitted a reinforcement learning model in which the value (V) guiding choice is a weighted mixture (w) of subtask-level (Q_ts_) and stimulus-level value (Q_a_), V = w·Q_ts_ + (1−w)·Q_a_, with the weight w estimated per participant and per condition (adapted from Eckstein and Collins, 2020). (A) Box plot showing participants had significantly higher accuracy for block compared to interleaved condition (43.5% vs 36.4%; z = 5.493 and p<0.001; Wilcoxon signed rank test; mean±sem). The solid dots represent individual participants’ data; grey lines connect the participants’ mean accuracy between conditions. The inset bar chart shows behavioural results from the MEG experiment, again showing higher accuracy in the block compared to interleaved condition (z = 1.943 and p= 0.0520; Wilcoxon signed rank test; mean±sem). All accuracies exceed chance level (theoretical baseline chance=0.042, 1/24 possible orders). (B) Fitted subtask-level value weight w by condition. Violin plot shows the across-participant distribution of w in block and interleaved conditions, with the horizontal marker denoting the mean. w was higher in the block than the interleaved condition (0.387 vs 0.228; Wilcoxon signed-rank p = 0.00791, dz = 0.24), indicating greater reliance on subtask-level value when a learned subtask structure was reusable. (C) Model predictive accuracy, exp(mean per-decision log-likelihood), in each condition. This metric informs how well the model explains the data. The dashed line marks chance level for a random four-stimulus ranking (0.347). Predictive accuracies in both conditions are above chance level (Wilcoxon signed-rank p < 0.001, dz = 1.24 for block and dz = 1.05 for interleaved). (D) Scatter plot showing parameter recovery of weighted mixture *w* (i.e. reliance on subtask value). Simulating rankings from known parameters on the empirical trial structure, and refitting them 1,000 times (each dot is one time), revealed a moderate Spearman rank correlation (rho = 0.408, p<0.001), suggesting successful recovery of the parameter. Dashed line is a diagonal perfect fit as a reference. (E) Forward simulation: the fitted model was run on participants’ recorded learning sequences, and the probability that it would produce the correct ranking was averaged across trials (exact-ranking accuracy, dashed line reflects chance = 1/24). The model was more accurate in the block than the interleaved condition (46.6% vs 39.0%; Wilcoxon signed rank test, p<0.001, dz = 0.51), reproducing a key direction of the behavioural effect in (A). Error bars denote s.e.m. across participants.

We next asked whether this behavioural advantage scaled with the amount of reusable structure. For this we divided interleaved problems into those wherein one subtask ranking remained intact (one subtask reusable) and those in which no subtask was reusable, in effect providing three structures (the block condition structure and the two interleaved condition structures). Accuracy differed across the three structures (mean accuracy = 43.5%, 37.3%, and 35.1% for block, interleaved with one reusable subtask, and interleaved with no reusable subtask, respectively; chi2 = 30.11, p < 0.001 with Friedman test), an effect driven by better performance in the block than interleaved condition subtypes (one-reusable-subtask z = 4.110, no-reusable-subtask z = 5.385, both p < 0.001; Wilcoxon signed-rank tests). However, the two interleaved subtypes did not differ (z = 1.5924, p > 0.1). Thus, an accuracy benefit was specific to block, with partial reusability conferring no significant advantage. Hereafter, we focus on block and interleaved conditions.

To identify participants’ precise behavioural strategies, we next fitted a reinforcement learning (RL) model to the online data (3 participants were excluded because of legacy data compatibility issues, yielding a total of N=238 participants for modelling; see Methods). In the model, we followed the spirit of hierarchical task-set value, (*V = w·Q_ts_ + (1 - w)·Q_a_*), where value (*V)* guiding choice is a weighted mixture *w* of subtask-level value *Q_ts_* and stimulus-level value *Q_a_*, as adapted from a previously described hierarchical task-set value model (Eckstein and Collins, 2020).

In the task, values are learned when they are revealed after each subtask choice, via a Rescorla-Wagner rule. Both *Q_ts_* and *Q_a_* values are reset at the start of every trial as stimulus values are re-randomised at each trial. In effect the weight w (0 ≤ w ≤ 1) indexes the degree to which choice is governed by subtask-level rather than stimulus-level value: w = 0 recovers a purely stimulus-based strategy, and w = 1 a purely subtask-based one. Because participants produced a full ranking rather than a single choice, the planning-phase ranking was modelled with a Plackett–Luce likelihood, a sequence of softmax choices made without replacement over the stimuli not yet ranked, using a single inverse temperature β. Values were acquired during subtask learning, but only the planning ranking entered the likelihood, mirroring the task demand placed on participants. On a proportion of trials, rankings were produced at random rather than from learned value, with its frequency defined by a lapse rate (ε). Five parameters, including the stimulus-value learning rate (α_a_), subtask-value learning rate (α_ts_), and the three parameters previously introduced (β, w, and ε) were all free, estimated per participant and per condition by maximum likelihood (see Methods).

To ensure the weighted mixture model is legitimate, and to select the best model, we tested the reliability of model comparison tools. For the reliability test, we first performed model recovery using the parametric bootstrap cross-fitting method (PBCM), bootstrap likelihood ratio test (BLRT), Akaike Information Criterion corrected (AICc), and Bayesian Information Criterion (BIC) to select the best method for comparing models (Supplementary text and Table S1-S3). Then, we compared the mixture model against alternatives, namely subtask-only learning (w=1), stimulus-only learning (w=0), and models without a lapse rate (ε), using the selected method, PBCM. Based on the model comparison results (Supplementary text), we rejected the alternatives and therefore used this model for analyses hereafter.

Within the weighted mixture model framework, we hypothesised a greater use of subtask-level value in the block condition than in the interleaved condition. Confirming this hypothesis, we found a significantly higher w in the block than interleaved condition (0.387 vs 0.228; Wilcoxon signed-rank p = 0.00791, dz = 0.24; Fig. 2B). In addition, we found a higher subtask value learning rate (α_ts_; p=0.00504, dz=0.24) and choice stochasticity inverse temperature (β; p<0.001, dz=0.29) in the block, indicating participants learned subtask value faster and made choices more consistently with learned value in the block compared to the interleaved condition. Other parameters showed no significant differences between the two conditions (α_a_: p = 0.365, dz = −0.08; ε: p = 0.811, dz = 0.05).

We next tested the robustness of the model, asking whether it predicted human behaviour with above-chance probability, in both conditions, and whether the parameter *w* could be recovered in simulated data. Expressed as predictive accuracy, the exponentiated mean per-decision log-likelihood (i.e. geometric mean probability) reached 0.516 in the block and

0.489 in the interleaved condition, each exceeding 0.347 expected under a random ranking (both Wilcoxon signed-rank p < 0.001, dz = 1.24 for block and dz = 1.05 for interleaved; Fig. 2C). We confirmed recoverability of the weight parameter *w* by simulating model behaviour, and refitting models (1,000 synthetic datasets), across a whole range of parameters, on the same task sequence participants performed (Spearman ρ = 0.408, p < 0.001; Fig. 2D). Together with other recoverable parameters (Fig. S3), and a weak correlation between parameters (Fig. S4), this provides evidence the model can identify parameter changes and potentially associated cognitive mechanisms.

We next asked if the across-condition differences in *w* could reflect bias arising from the task structure. We implemented two parametric simulation cohorts as controls (N=200 each), setting different null distributions for *w*: the first cohort was simulated using each participant’s own mean fitted w across conditions; the second cohort was simulated using *w*=0, as 90 participants preferred the *w*=0 model (supplementary text). After simulation, we refitted *w* to examine differences in *w* (*Δw*) across block and interleaved conditions (Fig. S5). The comparison of the empirical and null *Δw* gave one-sided bootstraps p=0.0448 for both cohorts, providing reassurance against artefactual findings caused by bias.

We then tested whether this model could reproduce the observed behavioural accuracy difference across conditions. For each participant, we ran the model forward with their own fitted parameters on their empirical learning sequences, using the fitted choice rule to compute the probability of correct rankings for the 4-stimulus planning problems (see Methods). The model was more accurate in block than interleaved conditions (46.6% vs 39.0%; signed-rank p < 0.001, dz = 0.51; Fig. 2E), replicating an empirical behavioural effect (Fig. 2A; 43.5% vs 36.4%). Moreover, we also found that, within each condition, simulated behaviour using participant-specific parameters and empirical accuracy were positively correlated (Fig. S6; block: Spearman ρ = 0.95, interleaved: ρ = 0.96; all p < 0.001). This suggests the model captures individual differences in performance and not merely a group-level condition effect.

These behavioural findings, involving an accuracy advantage in the block condition (Fig. 2A), together with a greater weighting of subtask value in that same condition (Fig. 2B-E) are consistent with the suggestion participants solved the planning task by exploiting subtask structure knowledge, within a condition when it serves a task solution.

Based on these behavioural findings we next asked whether MEG data provide evidence for reuse of subtask structure, where we hypothesised preferential subtask reactivation during planning in a block compared to interleaved condition.

### Subtask representational reactivation during compositional planning

A behavioural finding that participants avail of learned subtask features predicts enhanced subtask neural reactivation during compositional planning, specifically when subtask information is task relevant. To capture stimulus and subtask representations, we trained two distinct sets of decoders (see methods for details). A stimulus decoder was trained on the functional localiser prior to the main experiment, and a subtask decoder was trained during the subtask learning phase (when “Subtask 1”, “Subtask 2”, or “Subtask 3” text was shown on the screen). Both decoders showed significant above-chance performance under a leave-one-out cross validation (Fig. S7 & S8). As both decoders were trained on separate localiser and subtask-learning phases, rather than on the block or interleaved test trials, their cross-validated accuracy is independent of test condition (block/interleaved) and cannot, by construction, bias a subsequent condition comparison.

We applied these decoders separately to the task planning period, calculating average decodability across this window (self-paced, 5-30s). To isolate planning-specific neural reinstatement from general mnemonic or perceptual activation, we computed a differential reactivation measure involving decodability of task-relevant minus task-irrelevant stimuli (or subtasks). Here, task-relevant stimuli (or subtasks) were those needed to solve the current four-stimulus planning problem, whereas task-irrelevant stimuli (or the subtask) were those not needed. In light of previous research suggesting temporal differences for distinct features (Dobs et al., 2019; Xu et al., 2024), we illustrate this differential reactivation for decoders trained at each time point (Fig. 3A). To test whether neural reactivation differed between block and interleaved conditions and whether this difference changed across trials, we built Linear mixed-effects (LME) models predicting decodability, averaged across the within-trial window and decoding training time, from condition (block/interleaved), trial number, and their interaction, with predictors standardised to the same scale of −0.5 to 0.5.

**Figure 3.**
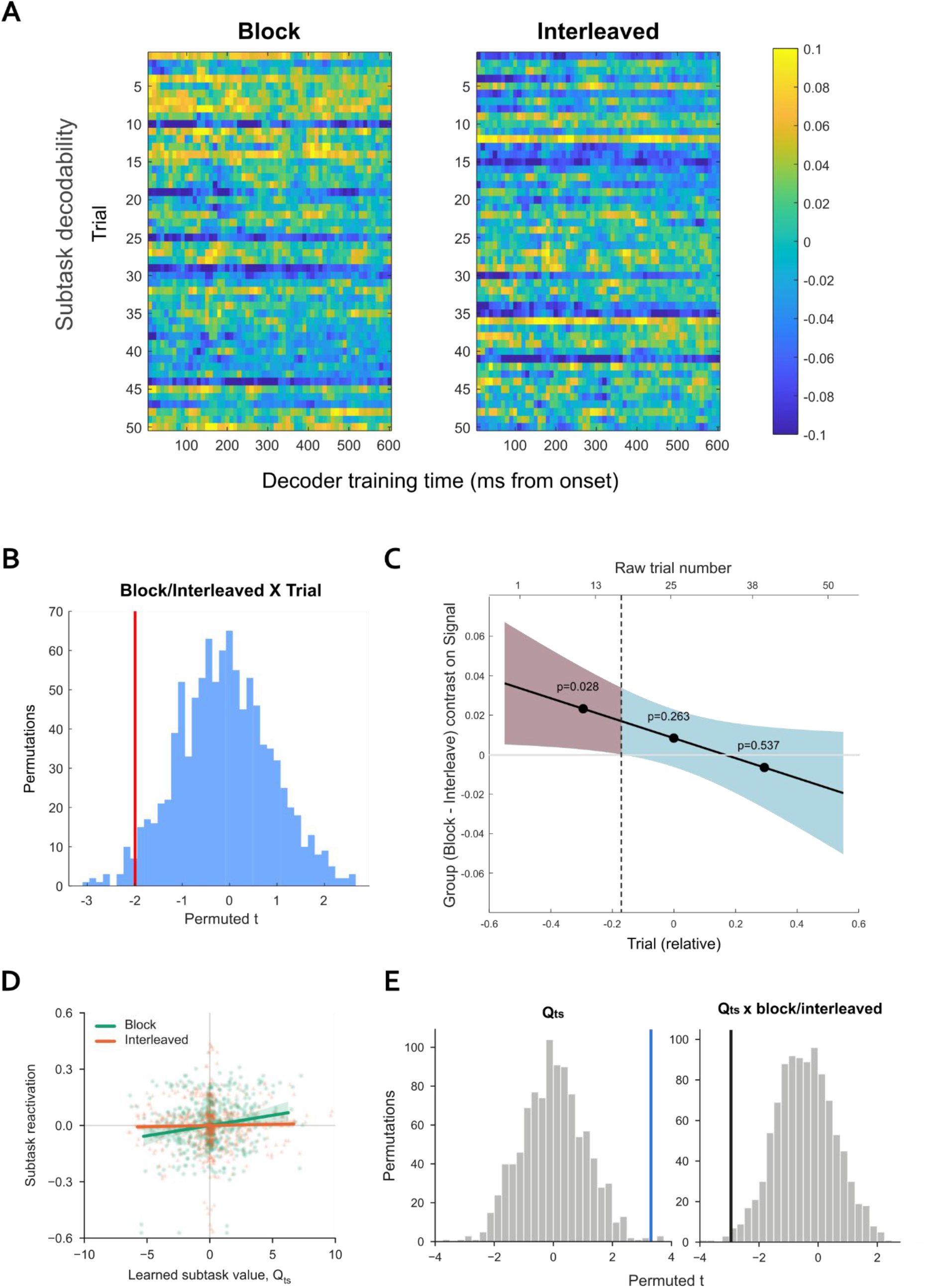
Reactivation decodability of subtask in block and interleaved trials. For stimulus decoding, we trained multivariate decoders for each stimulus (image) during the functional localiser task. For subtask decoding, we trained decoders during the problem-solving subtasks, when participants were presented with the word “Subtask 1”, “Subtask 2”, or “Subtask 3”. In a leave-one-out cross-validation these decoders performed significantly above chance (see Fig. S7 and S8). (A) Heatmap showing subtask reactivation decodability pattern for decoders trained at different time points and during different trials. Yellow indicates above-chance and blue indicates below-chance decodability. (B) Histogram showing the distribution of Condition × Trial t-statistics from the chunk-permutation null (N = 1,000) obtained by permuting condition labels at the level of consecutive same-condition runs within participant versus the observed t (red vertical line), with two-tailed p = 0.041. (C) Conditional block-minus-interleaved effect on subtask decodability across the session, as derived from the fitted LME. Shaded band, 95% CI; solid dots show simple-slopes tests at Trial = mean ± 1 SD (Aiken and West, 1991); dashed line shows Johnson–Neyman boundary (raw trial 17; Johnson and Neyman, 1936), with the red-shaded area showing region of significance at p<0.05 and the blue-shaded area showing non-significant (p>0.05) region. (D) Trial-wise subtask reactivation as a function of the mean learned value (*Qts*) of the two subtasks in the planning problem, shown separately for block (green) and interleaved (orange) trials. *Qts* was centred within participant and condition. Reactivation is shown as partial residuals after removing subtask-value difference, trial number and participant × condition means. Points represent individual trials, lines represent LME fits, and shaded bands represent 95% CIs from 2,000 participant-level bootstrap resamples. (E) Histogram showing permutation tests for relationship between reactivation and learned values (N = 1,000). Left, null distribution of the t statistic for *Qts*, obtained by shuffling *Qts* across trials within participant and condition, with two-tailed p = 0.005. Right, null distribution of the t statistic for the *Qts* × condition interaction, obtained by permuting condition labels at the level of consecutive same-condition runs within participant, with two-tailed p = 0.007. Vertical lines indicate the observed t statistics.

Subtask reactivation differed between conditions (block vs interleaved) in a manner that varied across trials, reflecting a condition × trial-number interaction. Thus, subtask decodability was greater in the block versus interleaved condition in early trials (quantified analyses below). A Linear mixed-effects (LME) model of per-trial reactivation (averaged across the within-trial decoding window and decoder training time) revealed a significant Condition × Trial interaction (β = −0.0505, two-tailed p = 0.0465), indicating enhanced subtask reactivation in the block relative to interleaved condition, an effect that attenuated over trials. To rule out a possibility this effect was driven by an autocorrelated blocked design, we conducted a chunk-permutation test, preserving each subject’s run-length structure (Fig. 3B; N = 1,000, shuffling the block/interleaved condition labels in chunks to avoid potential biases created by the poisson distribution structure). This analysis again revealed a significant effect (Condition × Trial with two-tailed p = 0.041).

To localise an interaction in time, we conducted simple slope (Aiken and West, 1991; Finley et al., 2024; Kelley et al., 2019) and Johnson–Neyman regions of significance analyses (Bauer and Curran, 2005; Johnson and Neyman, 1936; Katayama et al., 2026; Politte-Corn et al., 2025). The block-versus-interleaved contrast was reliable at trial 11 (β = 0.023, t = 2.20, p = 0.028) but not at trial 25 (p = 0.26) or trial 40 (p = 0.54), with the Johnson–Neyman boundary falling after trial 17, consistent with stronger subtask reactivation in early trials for the block condition (Fig. 3C). Interestingly, this selective reactivation in early session trials accords with rodent findings showing reactivation for repeated experiences systematically decreases over extended timescales (Chiossi et al., 2025; Mankin et al., 2012). By contrast, for stimulus representations, we found no differences in the LME models either for condition or its interaction with trial. The absence of an effect may reflect insufficient sensitivity of the stimulus decoder to capture condition-level differences or, alternatively, indicate that stimulus-level representations are comparably engaged across both conditions during planning.

While the main focus of this study is not to identify the neuroanatomical structures linked to compositional planning, we have run an exploratory source localisation analysis and found greater inferior frontal gyrus (IFG) source power during block planning than during interleaved planning (Fig. S9), suggesting potential involvement of IFG in planning with subtasks.

We next asked whether the strength of subtask reactivation reflected subtask learned values. For each planning trial, we calculated the planning mean learned value *Qts* of the two subtasks and tested whether higher value predicted stronger subtask reactivation, controlling for a value difference between the two subtasks. In an LME analysis, subtask reactivation strength increased as a function of mean subtask value (standardised β = 0.10, 95% CI [0.04, 0.15], p = 0.001), with positive participant-wise slopes in 20 of 25 participants (signed-rank p = 0.006, dz = 0.62). By contrast, the value difference between subtasks did not predict reactivation (β = −0.02, 95% CI [−0.08, 0.04], p = 0.47). Furthermore, this effect was present in the block condition alone (β = 0.19, 95% CI [0.10, 0.28], p < 0.001), but not in the interleaved condition (β = 0.02, 95% CI [−0.05, 0.10], p = 0.55; Fig. 3D), with participant-wise slopes showing the same pattern (block: signed-rank p = 0.039, dz = 0.43; interleaved: p = 0.56, dz = 0.10). Critically, a relationship between subtask value and reactivation differed significantly between block and interleaved conditions (*Qts* × condition: β = −0.17, 95% CI [−0.28, −0.06], p = 0.003). The robustness of this effect was evident in a finding that these results also held under permutation tests (Fig. 3E), where shuffling *Qts* across trials within participant and condition reaffirmed the relationship between subtask value and reactivation (two-sided p = 0.005, N = 1,000), while shuffling the block/interleaved condition labels in chunks supported an interaction effect (two-sided p = 0.007, N = 1,000). Thus, neural subtask reactivation scaled with learned subtask value when a subtask structure could be reused to solve a planning problem, paralleling a behavioural finding of greater weighting of subtask value in a block condition (Fig. 2B).

### Subtask representation transfer

Our task design afforded a further test of subtask representation and utilisation. Following the main experiment, participants were tasked to respond as to whether sequentially presented stimulus pairs derived from the same subtask or not (behavioural accuracy: 92.08% ± 2.87%, mean ± sem; p=0.0029, z=2.976, with a Wilcoxon signed-rank test against the majority baseline 80%, as we expect a “Yes” in only 1 out of 5 trials). Here we hypothesised enhanced subtask reactivation when presented stimuli align with this structure. Thus, if subtask representations are reutilised for problem solving, then this predicts they should also transfer to solve problems in novel contexts. To test this, we asked if subtask identity decoders (trained on the subtask word cue) generalise to the post-experiment probe question context (Fig. 4A). For this we applied decoders trained to distinguish the three learned subtasks (subtasks 1, 2, and 3) to MEG data corresponding with the point in time when participants viewed the second of the two probe images (−100 to 800ms after image onset). The justification for a focus on this time epoch is that it corresponds to when participants have all necessary information needed to determine whether the consecutively presented images belonged to the same subtask. Consistent with generalisation we found significant above-chance level decoding of subtask identity (cluster-based permutation test, N=1,000, FWE-corrected p<0.05), with peak decoding performance at 170ms post stimulus onset.

**Figure 4.**
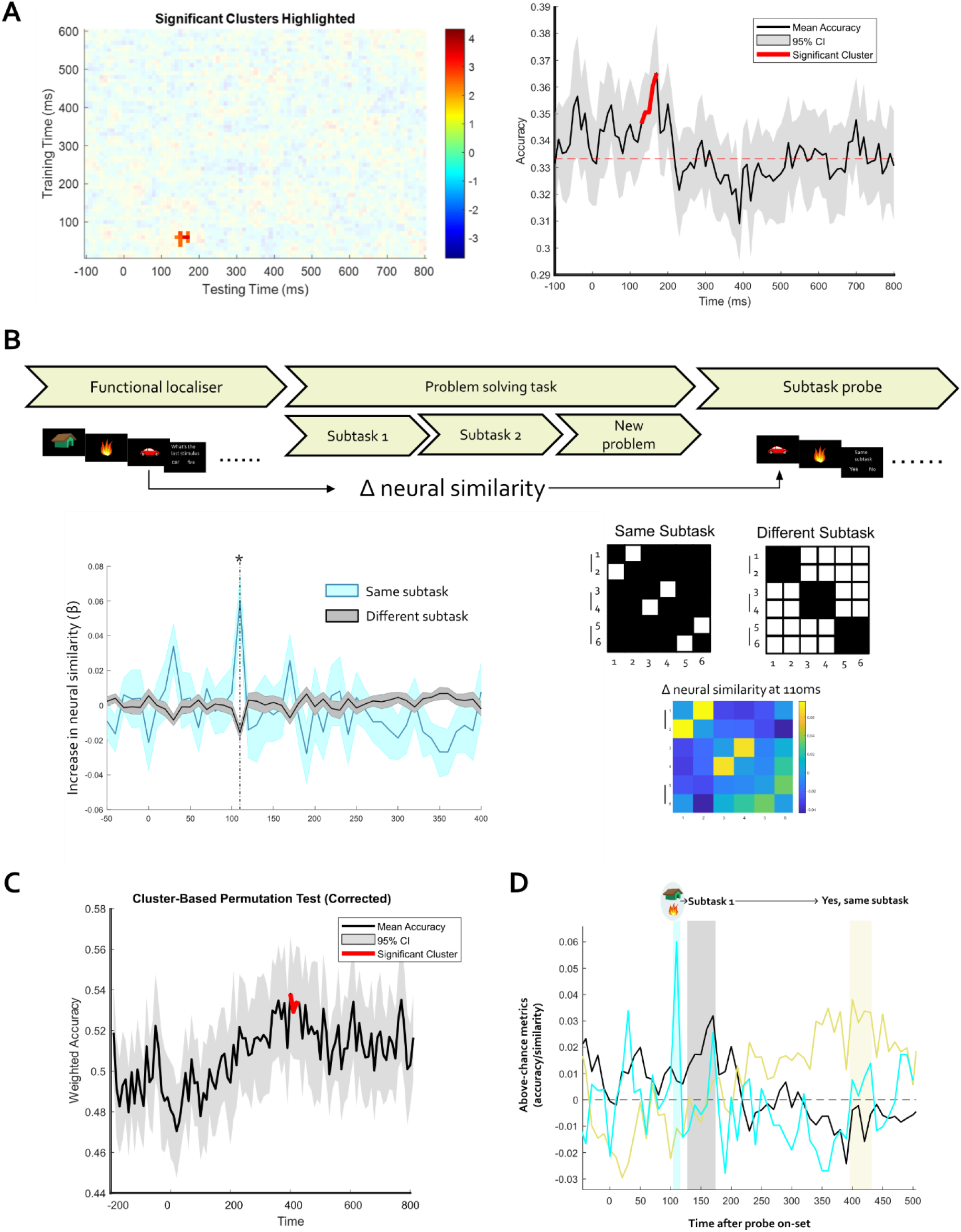
Subtask representation transfer. (A) Probe question subtask identity decoding. We asked whether trained subtask decoders (see Methods and Fig. 3) could decode subtask identity (1, 2, or 3) during the post-experiment probe question. We focused on when participants saw the second image, the time when they can ascertain if images do, or do not, belong to the same subtask. In a permutation test, shuffling labels (N=1,000), we found a significant positive cluster in the heatmap (left, significant cluster with FWE correction, thresholding at p<0.05 t-stats against 0). The right panel shows the performance of the decoder with the peak t-stats (60ms decoder training time; with the peak probe time at 170ms) in the heatmap. Dashed line represents chance level. (B) RSA analysis. This tests the hypothesis of task-induced neural similarity for two images belonging to the same subtask over the course of the experiment. We computed Pearson’s correlation between stimuli and used their differences as a metric for task-induced similarity change. We found a significant increase in neural similarity for stimuli (images) belonging to the same subtask (t = 4.162, p=0.0157, paired t-test corrected with Bonferroni correction, blue line), compared to stimuli pairs not from the same subtask (grey line). At the peak time point (110ms after stimulus onset), the top three increases in neural similarity were from representations within a subtask, as shown in the coloured matrices on the right. (C) MEG signals predict participants’ probe question responses (“Yes” or “No”). Due to the imbalanced trials, weighted accuracy was used, in which 0.5 is the chance level. Cluster-based permutation analysis (N=5,000, with FWE correction, shuffling probe question answer before training, thresholding at p<0.05 t-stats) tested if participants’ task responses, following onset of the second image, could be predicted from the neural signal. A significant cluster was found at 400-420ms. (D) Descriptive schematic summary of peak latencies for effects reported in panel A-C. In summary, a signature reflecting subtask similarity emerged at 110ms after probe image onset (blue shaded); a signature of subtask identity (grey shaded) showed a peak at 170ms; lastly, a signature related to probe question response appeared at around 400ms (yellow shaded). The shaded area surrounding the line represents the 95% confidence interval.

Previous studies report that with learning, stimulus representations adhering to the same task structure become more neurally aligned (Luettgau et al., 2025; Nour et al., 2021). Here we asked whether subtask learning was associated with increased neural similarity between images (stimuli) conditional on belonging to the same subtask. For this, we availed of a representational similarity analysis (RSA) and constructed a neural correlation matrix between stimuli from functional localiser data (prior to the main experiment) and again when participants were presented with individual images during the probe question task (after the main experiment; see Methods). Subtracting the two matrices yielded a [stimulus x stimulus] matrix of pre-to post-change in pairwise neural similarity. With six stimuli arranged into three two-stimulus subtasks, this yielded three within-subtask pairs and twelve between-subtask pairs (Fig. 4B matrices). As shown in Fig. 4B, a significant increase in neural similarity was evident for within-subtask stimulus pairs (at 110ms; mean differences = 0.0759, t = 4.162, p=0.0157, Bonferroni-corrected paired t-test) by comparison to between-subtask pairs. Notably, at this timepoint, the largest increases in Δ neural similarity matrix (Fig. 4 matrices) all correspond to the within-subtask pairs. In other words, increased within-subtask similarity was observed at a specific time point in a trial rather than across the entire period.

A reactivation of a subtask representation during the probe question led us to also ask whether the MEG data contained a neural signature of participants’ upcoming decisions. In a cluster-based permutation test, using weighted accuracy to account for trial imbalance (as a “Yes” was expected on only 1 in 5 trials), we asked whether participants’ dichotomous behavioural responses (“Yes”, same subtask; or “No”, different subtask) could be predicted from brain activity. We found probe responses (Yes/No) were indeed decodable (5,000 permutations, FWE-corrected p<0.05) from brain activity 400ms following onset of the second image (Fig. 4C).

The three probe-task effects peaked at different latencies: within-subtask similarity at 110 ms, subtask identity at 170 ms, and probe response at 400 ms (Fig. 4D). Qualitatively, this suggests subtask-related information is represented in a neural signal prior to a later probe response. However, because these latencies were estimated independently from separate analyses, this pattern should not be taken as evidence for a sequential processing cascade.

### Replay of subtask and stimulus during planning

Differential reactivation of subtask representations during planning led us, a final analysis, to ask if there was evidence for sequential reactivation (replay) of subtask and stimuli representations. To quantify sequenceness (sequential reactivation strength), we used temporally delayed linear modelling (TDLM; see Methods) (Liu et al., 2021a, 2019) with sequenceness defined in terms of ranking needed to solve the planning task on each trial. The subtask order was determined by the order of subtask identity appearing in the correct answer. For example, the correct target sequence of trial n in Figure 1A is fire->house->car->tree (subtask 1->subtask 2), while the correct target sequence of trial m is car->fire->tree->house (subtask 2->subtask 1).

Testing for subtask and stimulus replay during planning (across both conditions), revealed significant forward subtask replay at a 50-60ms time lag, exceeding a 95th percentile permutation threshold (Fig. 5A; with another later cluster at 230-290ms), with no evidence for significant forward replay for individual stimuli representations. For backward replay, we found significant effects for both subtask (Fig. 5B; with time lags of 70ms, 130ms, and 260-280ms) and stimulus sequences (with time lags of 90-100ms). We then tested for replay differences in block and interleaved conditions using cluster-based permutation tests, calculating replay sequenceness separately for the two conditions and comparing them (Fig. S10). We found marginally stronger forward subtask replay in the block condition than the interleaved condition (p=0.0779; FWE-corrected; Fig. S10A), and significantly stronger backward subtask replay in the interleaved condition compared to the block condition (p=0.0370; FWE-corrected; Fig. S10B). For stimulus replay, no significant between condition differences were evident (Fig. S10C and S10D; p>0.1; FWE-corrected).

**Figure 5.**
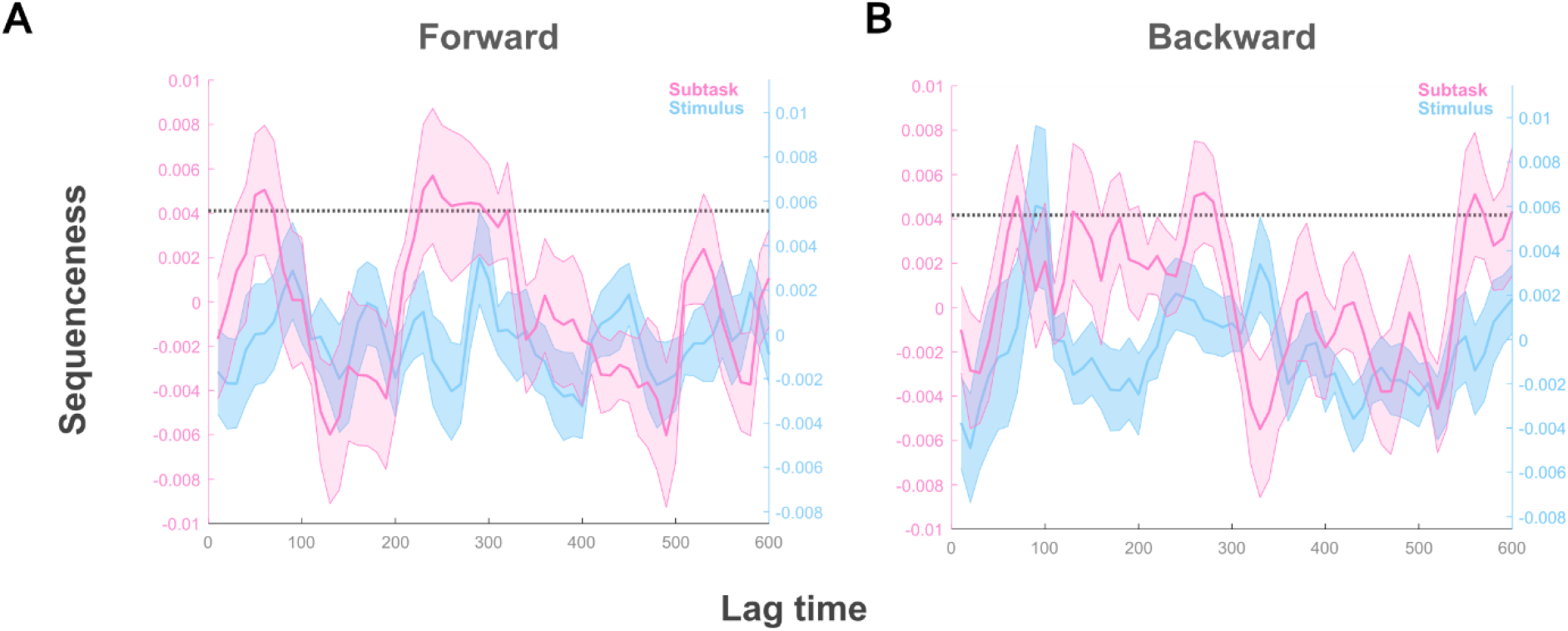
Stimuli and subtasks replay differently during planning. As in previous studies (Liu et al., 2019; Schwartenbeck et al., 2023), we used TDLM to index replay sequenceness (strength of replay) during planning. We used stimulus (image) and subtask decoders (see Methods for details), to find stimulus and subtask replay, respectively. (A) Significant subtask forward replay was seen at 50-60 ms time lags (together with another later time-lag at 230-290 ms). In contrast, we found no significant stimulus forward replay. Dashed line represents the significance threshold. (B) For backward replay, we found both significant stimulus (90-100 ms) and subtask replay (at 70ms, 130ms, and 260-280ms). Dashed line represents the significance threshold. The shaded area surrounding the line represents the 95% confidence interval. In addition, we also implemented a stratified replay analysis (Fig. S10), testing for differences in replay between block and interleaved conditions.

Behaviourally, participants performed better when tasked with high-to-low rankings compared to low-to-high rankings (Fig. S2), where in the latter there is incongruency with subtask ranking sequence presentation during learning (which is always high-to-low).

Previous studies, in rodents (Ólafsdóttir et al., 2017) and humans (Wimmer et al., 2023), report replay strength increases when a task becomes more demanding. Consistent with these observations, backward subtask replay was greater in low-to-high than in the high-to-low context (one-tailed p=0.002, FWE-corrected; Fig. S11), whereas no significant between-context differences were observed for other expressions of replay.

Together, these results are consistent with an expression of replay during planning not only for the individual stimuli, but also for subtask representations, where strength of replay is modulated by task context.

## Discussion

Our findings provide evidence that humans flexibly reuse learned subtask structure when it facilitates solving a complex problem. Behaviourally, participants were more accurate when previously learned subtask rankings could be directly exploited, with computational modelling supporting a greater reliance on subtask-level value information in this condition. At a neural level, task-relevant subtask representations were preferentially reactivated during planning, where strength of reactivation related to subtask value. Moreover, these subtask representations transferred to a novel probe context, and showed sequential forward and backward replay. The results provide convergent behavioural, computational and neural evidence that learned subtask representations are reinstated and redeployed during planning, providing novel insights that such reuse supports compositional problem solving.

The behavioural advantage conferred by reusable subtask structure suggests that participants do not solve each novel problem solely by reconstructing the values of its individual elements. Instead, the evidence suggests previously learned subtask level information contributes to subsequent planning, an interpretation supported by computational modelling showing greater weighting of subtask-level (*w*) information when this structure can be reused. A reuse of a learned structure provides a computationally efficient means of reducing planning demands by exploiting solutions to previously encountered subproblems. This supports the idea that reinforcement learning can operate over task-sets rather than individual stimulus-action pairs (Eckstein and Collins, 2020), where task-set values are redeployed when a new problem can be decomposed into already solved components. This reuse is particularly informative given values in human reinforcement learning are shaped by outcomes available in a local context (Bavard and Palminteri, 2023; Molinaro and Collins, 2023). The observed behavioural benefit from reusing learned subtasks implies that learned subtask values can be conserved across contexts to guide new problem solutions, an advantage evident in particular when the problem was composed of two learned subtasks.

This implies that behavioural gains depend on the availability of a sufficiently complete decomposition of a problem into reusable components, rather than scaling simply with the amount of previously learned structure.

Conceptually, subtask reuse bears more than a passing resemblance to dynamic programming. The core idea in the latter is that solutions to subproblems are stored and reused without needing to be recomputed. While our study did not test dynamic programming in a formal algorithmic sense, we de facto test a core computational principle that retaining intermediate solutions facilitates reduction in the complexity of future encountered problems (Bellman, 1966; Bellman and Dreyfus, 2015; Cormen et al., 2022). Analogously, during multi-step planning humans are adept at task decomposition using heuristics that reduce a complex task into its more discrete elements (Bein and Niv, 2025; Correa et al., 2023; Duncan et al., 2020; Eckstein and Collins, 2020; Franklin and Frank, 2018; Solway et al., 2014), with previous work indicating humans adhere to hierarchical and resource-rational variants of such a framework (Balaguer et al., 2016; Correa et al., 2025; Solway et al., 2014). Our behavioural and modelling results show that subtask-level values are utilised when subtask structures are beneficial for problem solving (Fig. 2B-E), a finding that may be relevant also for model-based control that involves computations over structured internal models (Daw et al., 2011; Moran et al., 2019).

Prior neuroimaging and electrophysiology studies have identified indices of hierarchical planning (Balaguer et al., 2016; Liang et al., 2022), compositional inference (Schwartenbeck et al., 2023), combining cognitive maps (Park et al., 2020), and multi-step reinforcement learning (Daw et al., 2011; Miller and Venditto, 2021). By demonstrating reactivation of higher-level learned subtask representations during planning, our work extends on previous findings showing exploitation of shared compositional elements in humans (Bernardi et al., 2020; Courellis et al., 2024; Lee et al., 2026) and animals (Tafazoli et al., 2026). Notable here is our finding that reactivation strength scaled with model-estimated subtask value, particularly in a block condition where computationally greater weight is assigned on subtask-level information, linking a neural signal to a key quantity expressed in choice.

Subtask representations were preferentially reactivated early in a scan session, when solutions were less familiar, but less strongly reactivated across repeated experience. A possible explanation for this is that early reactivation provides access to previously learned subtask representations for incorporation into a nascent planning policy, with this need reducing once a policy becomes established. Evidence consistent with this suggestion comes from reports of a similar decline in reactivation with repeated experience in rodents, specifically when they encounter previously explored environments during navigation tasks (Chiossi et al., 2025; Giri et al., 2019; Mankin et al., 2012).

Compositional generalisation is considered a hallmark of flexible intelligence in both humans and in AI (ARC Prize Foundation, 2026; Chollet, 2019; Lake et al., 2017; Lake and Baroni, 2023; Piray and Daw, 2025). Generalisation is thought to rely on access to representations that can be both shared and recombined across tasks, rather than bound to a single context (Ito et al., 2022; Mill and Cole, 2025; Schwartenbeck et al., 2023). Our results show subtask representations are deployed flexibly across contexts as compositional building blocks for planning as well as in the context of an entirely new task structure.

A methodological advantage of MEG is that it allows non-invasive access to human data at a millisecond-scale temporal resolution (Gross, 2019; Gwilliams and King, 2020; Sejnowski et al., 2014) that allows capture of the temporal unfolding of computations. Strikingly, we found decoders trained during the main experiment generalised to another context, consistent with a decoded subtask capturing a high-level representation. During a subtask transfer probe question, within-subtask similarity, subtask identity and decision-related signals peaked at around 110, 170 and 400 ms, respectively (Fig. 4D). While these latencies were estimated from separate analyses, this temporal ordering hints at a probable mechanism of how previously learned compositional representations are progressively engaged to solve a novel question, consistent with initial activation of a learned associative structure and a subsequent progression in support of a decision (Gwilliams and King, 2020).

While the sensitivity of MEG for detecting reactivated sequenceness in humans has been questioned (Kern et al., 2026), we found significant forward and backward sequenceness at the subtask level, in addition to replay of individual stimuli, consistent with replay operating over learned abstract subtasks rather than only sensory features alone. In addition, we also found context-dependent differences in the subtask replay. Relevant here is rodent work linking backward replay to value-driven learning (Ambrose et al., 2016; Diba and Buzsáki, 2007; Foster and Wilson, 2006; Ólafsdóttir et al., 2017) which is echoed in our finding of greater backward replay when stimulus-value associations provide the only route to a task solution (Fig. S10). Likewise, our results suggest stronger backward subtask replay when participants needed to reverse a ranking from high-to-low to low-to-high (Fig. S11), potentially reflecting increased task-demand (Ólafsdóttir et al., 2017; Wimmer et al., 2023) and/or replay of the original high-to-low sequence.

In an exploratory analysis, using MEG source localisation, we identified the left inferior frontal gyrus as a candidate substrate supporting subtask-structured planning (Fig. S9), with greater broadband source power (1-45 Hz) in the block compared to interleaved condition. Left inferior frontal gyrus (IFG) is implicated in hierarchical control across cognitive domains (Friederici, 2019; Jeon and Friederici, 2013; Koechlin and Jubault, 2006; Thompson-Schill et al., 1997), and greater engagement in a context where subtask utilization is beneficial is consistent with such control. Future work could examine more directly whether IFG signals exhibit a factorised geometry as predicted by theoretical accounts of multi-task learning (Wakhloo et al., 2026).

We acknowledge limitations to our study. First, our task involves a specific instance of compositionality (ordinal ranking within subtasks), and the generality of the mechanisms we detail to other forms of compositional problem-solving (e.g., spatial navigation, linguistic composition) remains to be established. Second, at best source localisation provides only an approximation to the likely neuroanatomical region linked to subtask utilisation (Haufe et al., 2011). Third, replay simulations have reported statistical challenges in replay detection using TDLM (Kern et al., 2026), which in our study may obscure the precise contribution played by neural replay signals. Last, our study does not provide causal evidence regarding how precisely reactivation contributes to compositional planning.

In summary, we provide behavioural, computational, and neural evidence that the human brain reactivates, replays, and transfers compositional subtask representations to solve complex problems. The results are a step toward a broader understanding of the neural basis of flexible compositional intelligence, a capability central to human cognition and AI development.

## Methods

### Participants for the online experiment

A total of 294 healthy participants based in the UK were recruited online with Prolific (https://www.prolific.co/), with 241 participants included in the final study after excluding participants who variously reported ‘didn’t know what to do’, gave up, had below chance-level accuracy, or didn’t finish the study. For the modelling, three participants were further excluded because of legacy data compatibility issues. Therefore, a total of 238 participants were used for modelling. All participants reported absence of a history of mental health and/or neurological conditions. As in previous research (Luettgau et al., 2024), participants were recruited only if they reached an approval rate of 95% from all other previous online experiments on Prolific. Participants received, on average, a compensation of £10.19 for the one-hour online experiment. The study was approved by the UCL Research Ethics Committee (ethics number: 3451/001).

### Online task design

The online experiment implemented a ranking task design illustrated in Fig. 1A. In each of 30 trials, participants first learned to rank two visual stimuli (from high to low) in each of the three subtasks based on trial value, randomly assigned at each trial (from −10 to +10). Using a cursor, participants then clicked on images sequentially to indicate their rankings within a 4s allowed window. Across clicks, feedback was provided regarding the value of each image (shown on screen for 1s). After that, they were informed if the ranking was correct or not (for 2s).

Once participants correctly answered a ranking twice in a row, they entered into the next subtask and this continued up until they learned a set of three subtask rankings. In their first attempt, as participants did not know the value of any stimuli this was not considered for a “twice-in-a-row correct” learning criterion.

After learning three subtasks, participants were tasked to solve a complex 4-stimuli ranking problem, comprised of two randomly selected subtasks from the set of three. For the planning task, we first showed the participants the 4 selected images (2s) and the ranking rule (high-to-low or low-to-high; for 2s). Unbeknown to participants, we imposed two conditions: in a block condition, within-subtask ranking (for example, clicking “Fire” first and then “House”) could be reused directly; in an interleaved condition, they need to rely solely on individual value-image associations, as another stimulus rank was interposed between (for example, “Tree” with value of 3, which is in between “Fire” with value of 5 and “House” with the value of 2).

Subjects then entered a planning period of variable duration of 5-30 seconds (self-paced), and committed to a choice (click the 4 images in correct sequential order according to their values, within 10s). Outcomes (correct or wrong) were provided and a 10-pence bonus given for each of the correct answers on the 4-stimuli ranking task. To test whether staying in the same (block/interleaved) condition influences behaviour, we split the participants randomly into 5 categories during online recruitment, with the probability of changing condition following different Poisson distributions (expectation λ=1, λ=5, λ=10, only block, and only interleaved), thereby having 47-50 participants for each of these 5 categories.

### Computational models of behaviour

To characterise strategies underlying behaviour, we fitted a reinforcement learning (RL) model wherein choice is considered to be driven by a weighted mixture of subtask and stimulus-level value. The model shares the hierarchical scheme of Eckstein and Collins (2020), maintaining value at two levels of abstraction, but departs from it in three aspects as dictated by our task. First, subtask identity was explicitly cued rather than being a latent context to be inferred, so no task-set inference is required; the two levels instead compete through a single free weight. Second, participants produced a full ranking of four stimuli on each planning trial rather than a single categorical choice, such that a single-action softmax is replaced by a Plackett–Luce ranking likelihood. Third, stimulus values were re-randomised on every trial (−10 to +10) such that they cannot accumulate across the session. Therefore, all Q-values are reset to 0 at the start of each trial and learned within-trial.

### Model

The model maintains stimulus-level values Q_a_ (one entry per stimulus, six in total) and subtask-level values Q_ts_ (one entry per subtask, three in total). Ranking the two stimuli of a subtask constituted two successive choices, with each choice treated as a separate learning event. On each event, the value of the chosen stimulus and the value of its subtask were both updated towards r, the point value of the chosen stimulus revealed on that trial (−10 to +10), via a Rescorla-Wagner rule:

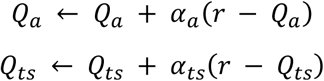

where α_a_ and α_ts_ are the stimulus-and subtask-level learning rates. These updates ran on every subtask learning trial, but only the 4-stimulus planning rankings (not the subtask choices themselves) were used for the likelihood evaluation. At the planning stage, the overall value of a stimulus (weighted mixture w of the two Q) enters the choice rule:

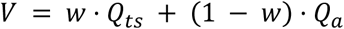

with w ∈ [0, 1]. At w = 0 the model is purely stimulus-based (equivalent to a flat model); at w = 1 it is purely subtask-based. Because Q_ts_ is shared by both stimuli of a subtask, increasing w shifts whole subtasks relative to one another without altering the order within a subtask.

*Action selection.* Choice probability P at a single rank position followed a softmax over the available stimuli:

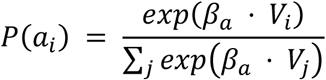

where β is a single inverse temperature capturing choice stochasticity. When participants ranked from lowest to highest value (as opposed to high-to-low), signs of value *V* were reversed before applying the softmax.

For 4-stimulus planning-task trials, in which participants produced a full ranking of stimuli (K = 4), the ranking likelihood was computed under a Plackett–Luce model. This treats a ranking as a sequence of choices without replacement: the probability of the full ranking is the product, across rank positions, of selecting by softmax the stimulus placed at that position from the stimuli not yet ranked. For an observed ranking (a_1_, …, a_K_):

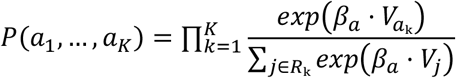

where R_k_ is the set of stimuli not yet placed by position k, so the highest-valued remaining stimulus is most likely to be placed next. On a proportion of trials, the ranking was produced at random rather than from the values, at a rate controlled by a lapse rate parameter ε following P(ranking) = (1 - ε) * P(ranking | value) + ε / 4!.

### Model fitting and comparison

The log-likelihood for each trial *t* was the Plackett–Luce log-probability of the observed ranking, LL_t_ = log P (a₁, …, a_K_), including the lapse. The value-based component, log P(a_1_, …, a_K | value) is summed over the K - 1 ranked positions (the final position is determined):

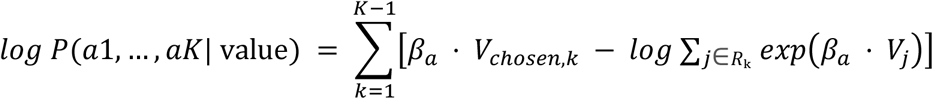

where V_chosen,k_ is the value of the stimulus placed at position k. The five free parameters (αa, αts, β, w, ε) were estimated separately for each participant and each condition by maximising the summed log-likelihood over that condition’s planning trials, using L-BFGS-B (scipy.optimize) with five random initialisations and retaining the best-fitting solution. Parameters were bounded at αa, αts ∈ [0.01, 0.99], β ∈ [0.1, 20], w ∈ [0, 1], ε ∈ [0, 0.5].

Fitted values of w were compared between conditions with a Wilcoxon signed-rank test (reported with dz).

### Predictive accuracy

To report model fit on an interpretable scale, we computed predictive accuracy as the exponentiated mean per-decision log-likelihood:

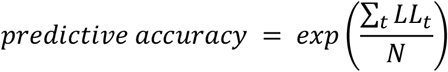

where N is the total number of ranking decisions (K - 1 = 3 per four-stimulus ranking). This is the geometric mean probability the model assigned to participants’ actual ranking decisions. The corresponding chance level, for a model with no preference, is the geometric mean of 1/4, 1/3 and 1/2, that is exp([ log(1/4) + log(1/3) + log(1/2)] / 3) = 0.347.

### Forward simulation of ranking accuracy

To test whether the fitted models reproduced the behavioural accuracy difference between conditions, we ran the model forward on the empirical task sequences (the 238 sets used in the online experiments). For each problem, Q-values were initialised to zero and updated on the participant’s actual subtask choices in the recorded presentation order, using the revealed stimulus values as reward, so that the block and interleaved orderings entered the simulation directly. After the subtask phase, we computed the 4-stimuli values V = w·Q_ts_ + (1−w)·Q_a_, using that participant’s fitted parameters. Accuracy was computed as the probability that the fitted choice rule (including the lapse) would produce the correct ranking, averaged across planning trials. This exact-ranking accuracy is comparable with the behavioural data as they are on the same scale (chance = 1/24). Block and interleaved conditions were compared with a signed-rank test.

### Parameter recovery

Because our inference rests on comparing a fitted parameter across conditions, we verified that parameters were recoverable. We drew 1,000 parameter sets at random from within the fitting bounds, simulated rankings from the model on the empirical trial structure (sampling each ranking from the Plackett–Luce policy), and refitted the model to these synthetic data using the exact same procedure. Recovery was quantified by a Spearman rank correlation between true and recovered values, robust to the compression that arises as parameters approach their bounds.

### Participants for the MEG experiment

Twenty-five healthy participants (15 female; age: 25.12 ± 4.38, mean ± std, range 18-34) were recruited for the MEG experiment. All participants reported normal or corrected-to-normal vision, with no history of any neurological or psychiatric disorders. The study was approved by the UCL Research Ethics Committee (ethics number: 14261/002). Participants gave written informed consent and received a base payment of £23.4 and an additional performance-based bonus up to £23.4.

### MEG task design

The MEG session comprised three phases: a functional localiser, a main experiment, and a final subtask probe phase.

### Functional localizer

Prior to the main experiment, each of six images was presented in pseudorandom order. Each image was presented 50 times, comprising 300 trials in total. Each image appeared on the screen for 1 second. In 20% of trials, there was a probe question, in which participants were tasked to press a button to indicate which stimulus had appeared on the previous trial. They responded with a controller, indicating left or right, where one of the two is the correct answer. A 2-pence bonus was awarded for each correct answer. This task was designed to train stimulus-specific decoders, following procedures established in previous work (Liu et al., 2019; Nour et al., 2021; Schwartenbeck et al., 2023).

### Main experiment

The main experiment followed the same design as the online experiment but comprised 50 trials with a Poisson distribution (change of condition λ=10). For each trial, participants first learned a 2-stimuli ranking (stimuli images randomised across participants) for three subtasks, with each requiring two consecutive correct responses before proceeding (the first attempt was not counted). They then used what they learned to solve a complex 4-stimuli ranking problem. All rankings were entered by sequentially pressing buttons on the controller matching the positions of the stimuli on the screen. A 30-pence bonus was awarded for each correct complex 4-image ranking. During planning, participants were given a time period of 5 to 30 seconds (self-paced), the primary window for MEG analyses of reactivation and replay.

### Subtask probe

After the main experiment, two images were presented sequentially on each trial, with participants tasked to indicate whether these images belonged to the same subtask or not. Each of the six stimuli was paired with each of the other five: one stimulus was its subtask-mate (a within-subtask pair), and the remaining four belonged to the other two subtasks (between-subtask pairs). The four between-subtask pairings of a stimulus were averaged into a single between-subtask estimate. Each stimulus was presented 50 times, with 6 stimuli, comprising a total of 150 trials (300 stimuli shown, with each trial containing two stimuli). For each correct answer, a bonus was given. To discourage participants from defaulting to “not the same subtask” (given this is the majority), a 4-pence bonus was given for a correct “same subtask” trial, while a 1-pence bonus was given for a correct “different subtask” trial.

### MEG data acquisition and preprocessing

MEG data were recorded using a 275-channel axial gradiometer scanner (CTF Omega from VSM MedTech) at a 1200Hz sampling rate at the Wellcome Centre for Human Neuroimaging, UCL. Three sensors were deactivated across all participants due to excessive noise. Participants were instructed to sit upright in the scanner and report answers using a scanner compatible controller (button box).

We followed a preprocessing protocol aligned with recent publications (Liu et al., 2019; Luettgau et al., 2025; Nour et al., 2021; Schwartenbeck et al., 2023). Data were resampled from 1200Hz to 100Hz to improve signal-to-noise ratio and high-pass filtered at 0.5Hz (with a first-order IIR filter) to remove slow drift. Independent component analysis (ICA) (FastICA, http://research.ics.aalto.fi/ica/fastica) was used to decompose sensor data into 150 temporally independent components. Artefactual components (e.g., eye blink and heartbeat) were automatically annotated and removed with classification based on spatial topography, high kurtosis and frequency spectrum. MEG triggers were realigned to photodiode signals recorded simultaneously with stimulus presentation to ensure the neural time series aligned precisely with behaviour. Different MEG sensors were excluded for each participant based on individual noise level (identified as noisy consistently throughout the task). All MEG analyses were performed based on filtered, cleaned MEG signals.

### Decoding analyses

*Stimulus Decoders.* We followed established procedures to train stimulus decoders (Liu et al., 2021b, 2019; Nour et al., 2021). Stimulus (image) specific decoders were trained on MEG sensor data from a functional localiser. At each time point post-stimulus onset, L1-regularised (LASSO) logistic regression classifiers were trained on the MEG sensor signals (channels as features) to discriminate between the six stimuli. In line with previous studies (Kurth-Nelson et al., 2016; Schwartenbeck et al., 2023), the LASSO penalty was fixed to λ=0.006. The decoder performance was benchmarked using a leave-one-set-out cross-validation (each fold left out 6 stimuli, one from each stimulus) procedure (Fig. S7). The trained classifiers were then applied to the MEG data during the planning.

### Subtask decoders

Similar to the stimulus decoder, we trained subtask identity decoders on MEG sensor data at the onset of a subtask word cue (“Subtask 1”, “Subtask 2”, “Subtask 3” shown on the screen) that preceded the subtask learning phase. LASSO logistic regression classifiers were trained to distinguish the three subtask identities, and the leave-one-set-out cross-validation results are reported in Fig. S8. We subsequently applied these classifiers to the MEG data during the planning task stage.

### Reactivation analysis

For each participant and trial, we quantified reactivation of each decoder (stimulus or subtask) during the planning phase. At every 10 ms of the planning phase, we computed decodability as the decoder’s predicted probability for the relevant state. To isolate task-relevant reactivation, we took a differential measure: decodability of stimuli (or subtasks) required on the current trial minus those not required, such that positive values index preferential reactivation of planning-relevant content. To ensure the result did not depend on a single feature (Al Roumi et al., 2021; Demarchi et al., 2019; Dobs et al., 2019; Xu et al., 2024), we trained decoders at multiple time points across the training epoch and averaged the differential decodability across these training time points and across the within-trial planning window, yielding one reactivation value per trial (for illustration, Fig. 3A shows this pattern before averaging across decoding time). This differential decodability was then entered into Linear mixed-effects (LME; *fitlme* in MATLAB) models to calculate how different predictors contribute to reactivation (with standardised beta estimates; all predictors were scaled to −0.5 to 0.5 before modelling), following the structure below:

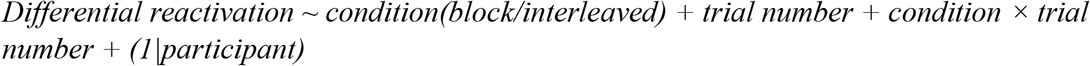

In addition to the standard LME fixed effects t-tests, we also assessed the statistical inference via a chunk-permutation test (N = 1,000, shuffling block/interleaved labels chunk-wise rather than per trial, so the temporal autocorrelation of the Poisson condition schedule is preserved).

*Relationship between learned subtask value and subtask reactivation*. For each planning trial, we reconstructed the learned value of each subtask (*Qts*), from the five-parameter model using each participant’s fitted parameters. We then calculated the mean (*Qts*) and absolute difference (|*ΔQts*|) of the values of the two subtasks in the problem. Both were mean-centred within participant and condition, such that only trial-to-trial variation entered the model, and z-scored across trials. Reactivation and trial number were also z-scored, yielding standardised beta estimates; condition was coded −0.5 (block) and 0.5 (interleaved). We fitted:

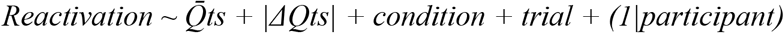

The same model, without the condition term, was fitted separately to block and interleaved trials. The difference between conditions was tested with:

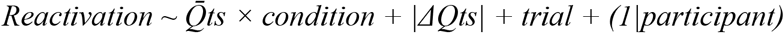

For permutation tests, *Qts* was shuffled across trials within participant and condition to test its main effect, while condition labels were shuffled across consecutive same-condition runs within participant to test the interaction (1,000 permutations each).

### Probe response decoding

Besides decoding the planning task, at the end of the experiment, we also assessed whether we could decode subtask identity as well as the participants’ response to the subtask probe question.

We used the same subtask decoders to decode subtask identity during the probe question phase, when participants were presented with the second image of the probe question and determined if the two images came from the same subtask (Fig. 4A). We used a cluster-based permutation test shuffling the labels of subtasks (N=1,000) to determine if their identity is decodable during this period.

Finally, we asked whether neural signals reflected the probe question decision, and when these signals emerge relative to subtask representations. By training decoders with MEG sensor data during a probe question, we asked if we could decode the participants’ answers (“Yes” same subtask; or “No” different subtask). We used LASSO logistic regression classifiers to predict the action of the participants. Weighted accuracy (5-fold cross validation) with cluster-based permutation analysis (N = 5,000, FWE-corrected, shuffling the answer labels) were used to determine if actual choice was decodable when participants viewed the second image (Fig. 4C).

### Representational similarity analysis (RSA)

As in previous studies (Luettgau et al., 2025; Nour et al., 2021), we tested whether learning in the main experiment increased the neural similarity of stimuli belonging to the same subtask. We computed the time-resolved RSA on MEG data from both the functional localiser and the subtask probe. At each time point, for each pair of stimuli (any pair, including both same and different subtasks), we computed the Pearson correlation between their MEG sensor signals. We calculated neural similarity differences (probe minus localiser) to ask whether within-subtask stimulus pairs (two stimuli associated with the same subtask; e.g. fire and house in Fig. 1) show enhanced similarity compared to between-subtask stimulus pairs (two stimuli associated with different subtasks; e.g., fire and tree in Fig. 1). Statistical significance was assessed via the Bonferroni-corrected (45 time points from −50ms to 400ms) paired t-test.

### Replay sequences quantification

To index replay sequenceness (sequential reactivation), we used temporally delayed linear modelling (TDLM) (Liu et al., 2021a, 2019). First, trained decoders with peak cross-validation accuracy (stimulus: 160ms, subtask: 180ms after stimulus onset) were applied to the MEG data during the planning phase, yielding a time-resolved reactivation (decodability) for each state (stimulus or subtask identity) at each time point. In effect, this transforms sensor data to decoded state space data. Next, we estimated its temporal structure, where for each time lag Δt (in steps of 10 ms, from 10 to 600 ms), the decoded state reactivation matrix at time t was regressed onto the same matrix shifted by Δt. This procedure yielded a transition matrix β(Δt), which was then regressed simultaneously onto four transition matrices: forward and backward transitions, defined according to the correct stimulus/subtask ranking for the planning task; an identity matrix capturing self-transitions (autocorrelation); and a constant matrix capturing non-specific co-activation across states. The regression weights for the forward and backward matrices provide a scalar measure of sequenceness at each time lag.

To assess significance, we permuted the state (stimulus/subtask) labels (N=1,000) and generated a null distribution of sequenceness at each time lag. Since there were limited possible permutations for subtask transitions, for each participant, we oversampled (bootstrapped using *bootstrp* in MATLAB) the shuffled subtask sequenceness to 1,000 data points, according to the distribution. To control for multiple comparisons, we set the significance threshold of replay as the 95^th^ percentile of sequenceness across all time lags (maximum across lags within each permutation) in the shuffled data.

To compare stimulus and subtask replay across block and interleaved conditions, we computed the sequenceness separately for block and interleaved trials. We then tested for a condition difference using cluster-based permutation analyses: first, time points where the t-statistic for the condition difference exceeded p = 0.05 were identified, and then the cluster mass was computed. Specifically, we used a two-tailed t-test, testing the differences between block and interleaved forward/backward replay. Significance was assessed against 1,000 permutations of the block/interleaved labels, with family-wise error (FWE) correction.

Similarly, we compared the low-to-high ranking situation to the high-to-low ranking situation in a similar cluster-based permutation analysis. Previous research (Mattar and Daw, 2018) has proposed that replay strength increases along with task difficulty. Therefore, in a one-tailed t-test, we tested the low-to-high>high-to-low situation, as participants need to flip the ranking over for the low-to-high situation.

### MEG Source Localisation

We performed source reconstruction using OHBA Software Library (OSL) and SPM12. A scalar linearly constrained minimum variance (LCMV) beamformer (Van Veen et al., 1997) was used to reconstruct epoched MEG data (0–5 s post-planning onset, band-pass filtered 1– 45 Hz) (Wimmer et al., 2020), to a volumetric grid in MNI space, with 5 mm isotropic spacing. The sensor covariance matrix was estimated over the planning window using broadband power, across all trials within each session, with PCA dimensionality reduction (50 components) for regularisation (Woolrich et al., 2011). For each trial, we computed broadband source power as the temporal variance of the reconstructed time course at each grid location. To remove a depth bias, we computed a normalised contrast per participant: (P_block_ - P_interleaved_) / P_grandmean_, where P_grandmean_ is the mean power across all trials from both conditions. One participant was excluded due to a global power imbalance between conditions (>3 SD from the group mean), leaving N = 24.

At a group level, participant contrast maps were entered into a one-sample permutation test (5,000 sign-flip permutations). No voxels survived whole-brain threshold-free cluster enhancement (TFCE) correction (FWE-corrected p < 0.05). Given an a priori hypothesis regarding left IFG in hierarchical planning (Koechlin and Jubault, 2006; Thompson-Schill et al., 1997), we performed a small-volume correction within an anatomical mask of left IFG (pars opercularis, pars triangularis, and pars orbitalis in the AAL3 atlas), applying TFCE with 5,000 permutations (FWE-corrected p < 0.05).

## Acknowledgements

This work is supported by the Max Planck UCL Centre, a joint initiative supported by the Max Planck Society and UCL. We acknowledge funding from the Max Planck Society (549771-D.CON 177814) and a Wellcome Trust Investigator Award to R.J.D. (098362/Z/12/Z). Maria K. Eckstein contributed in an advisory role only.

## Declaration of generative AI and AI-assisted technologies in the writing process

During the preparation of this work, the authors used ChatGPT and Claude in order to check grammar and code. After using these tools, the authors reviewed and edited the content as needed and take full responsibility for the content of the published article.

## Supplementary figures

**Figure S1.**
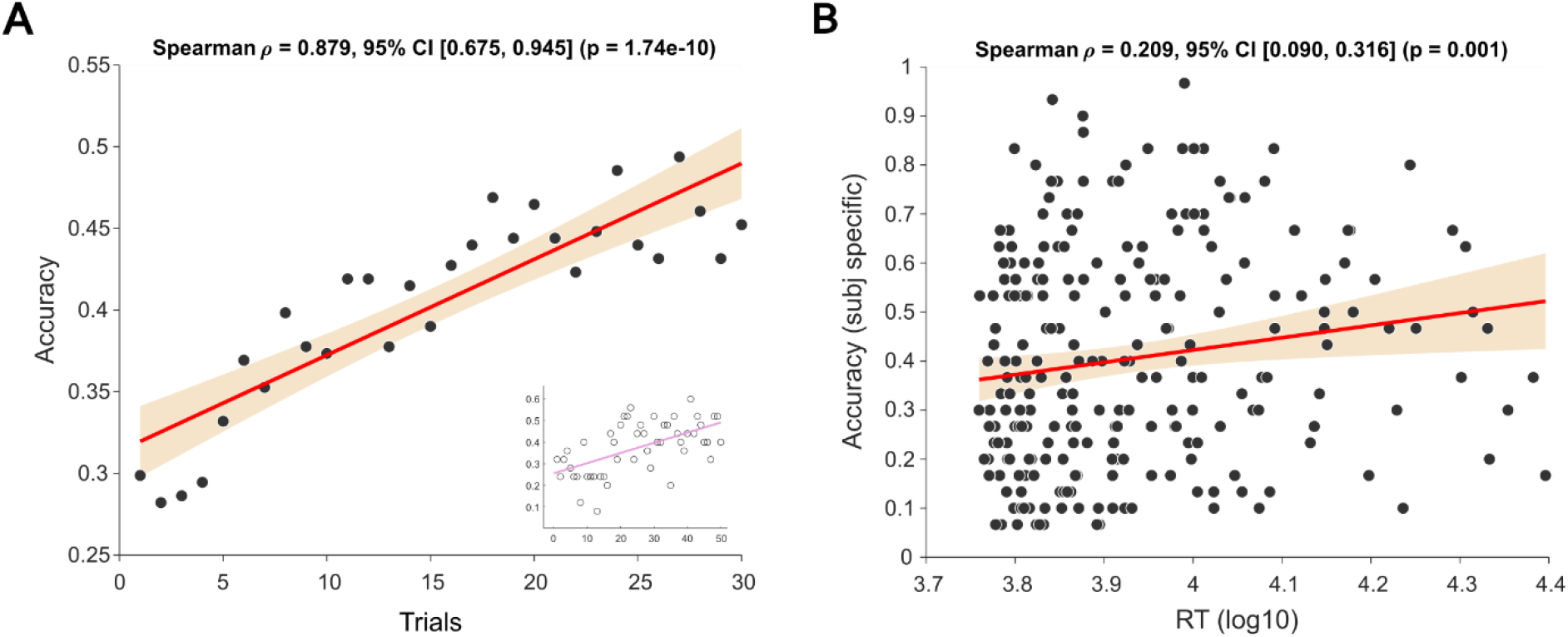
Behavioural features linked to task engagement. (A) Scatter plot and correlation analysis showing increased accuracy (behavioural performance) across trials. The orange shaded region shows the 95% confidence interval, and the red line shows the least-squares line of all dots. The inset scatter plot shows the consistent significant behavioural results in the MEG experiment (with the pink line representing the least-squares line). (B) Scatter plot and correlation analysis show increased accuracy for participants who manifest longer planning times. The orange shaded region shows the 95% confidence interval, and the red line shows the least-squares line of all dots.

**Figure S2.**
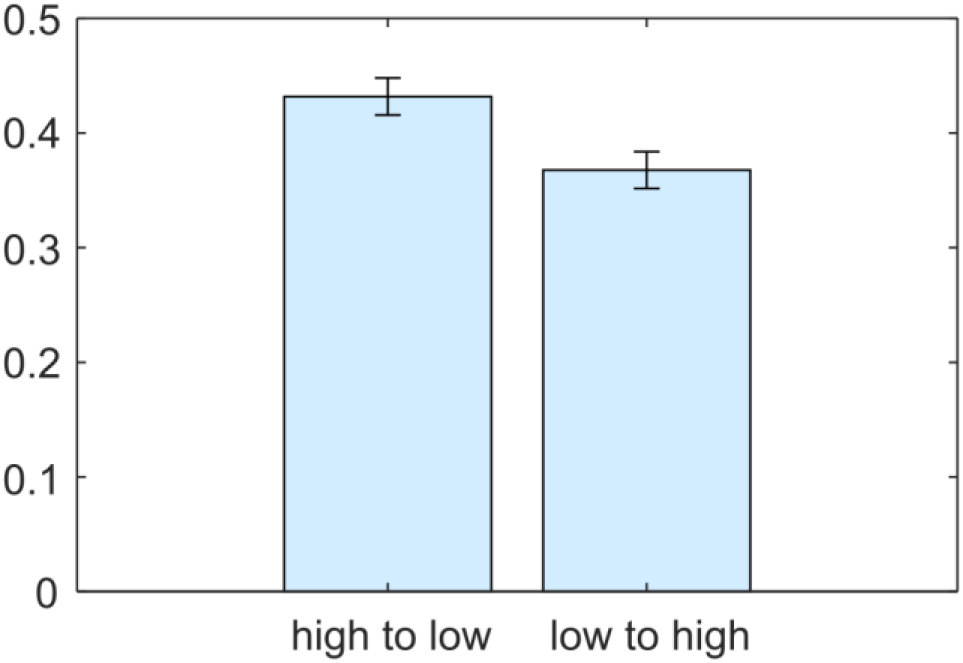
Accuracy with different ranking rules. Bar chart showing participants had significantly higher accuracy for the high-to-low than low-to-high situation (p= 3.950e-08; Wilcoxon signed rank test; N=241). Error bars represent mean ± sem.

**Figure S3.**
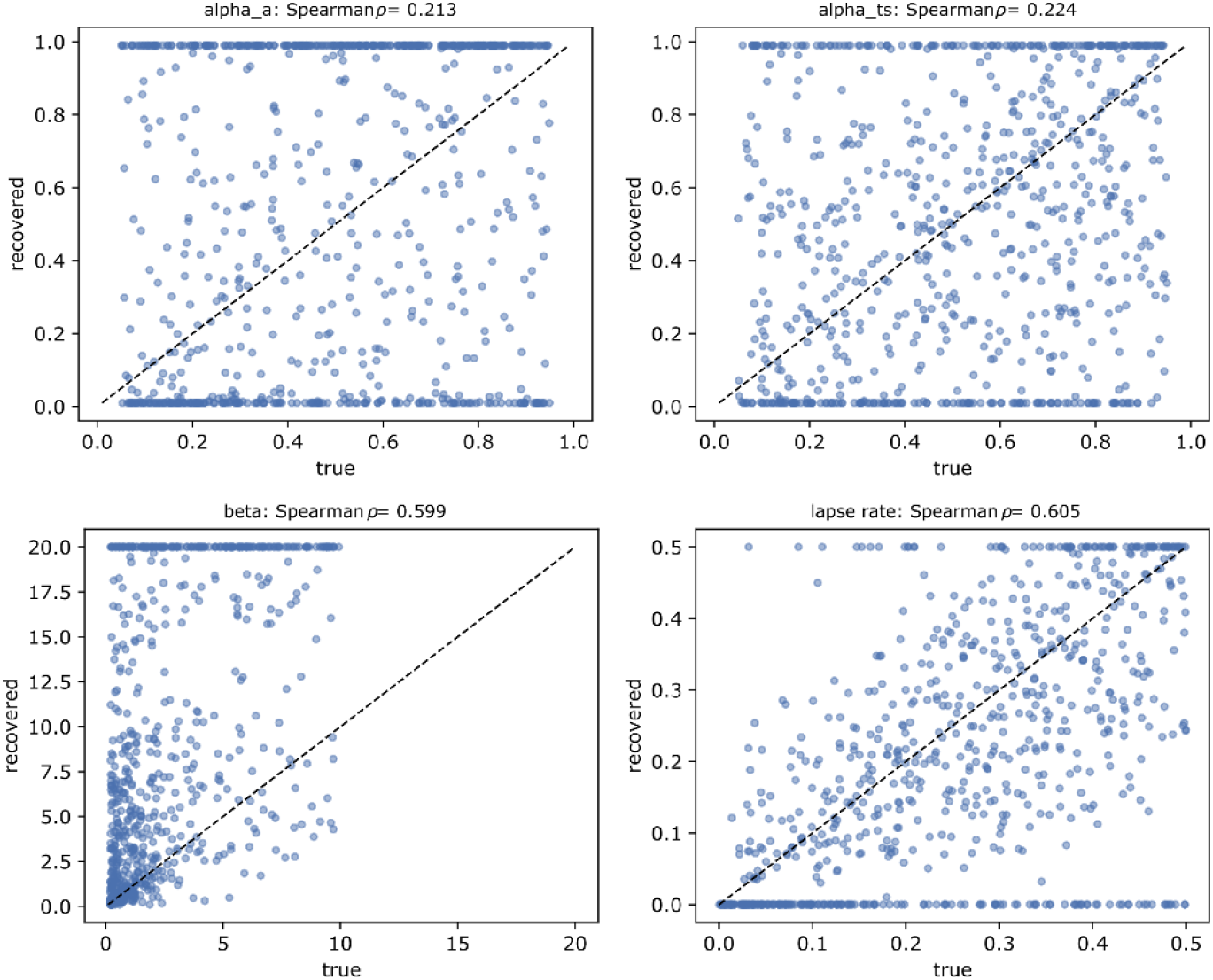
Parameter recovery of weighted mixture model. Scatter plots showing the recoverability of stimulus learning rate (α_a_, top left), subtask learning rate (α_ts_, top right), stochasticity (β, bottom left), and lapse rate (ε, bottom right) parameters. Each dot represents one simulation. The recovery of the mixture parameter w is shown in Fig. 2D.

**Figure S4.**
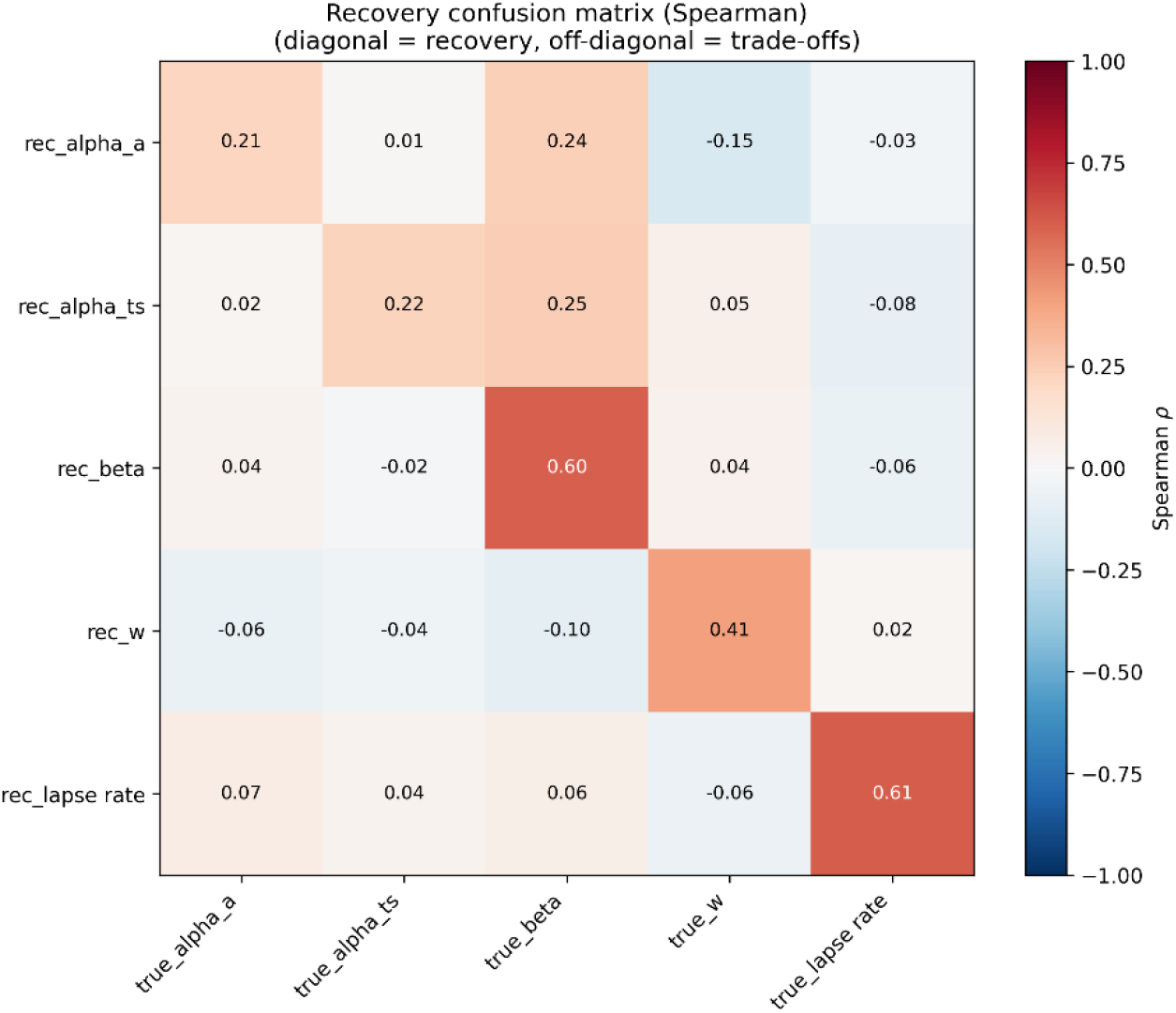
Parameter recovery confusion matrix. Confusion matrix showing prominent correlations come from the recovery (diagonal) but less so from the trade-offs (off-diagonal).

**Figure S5.**
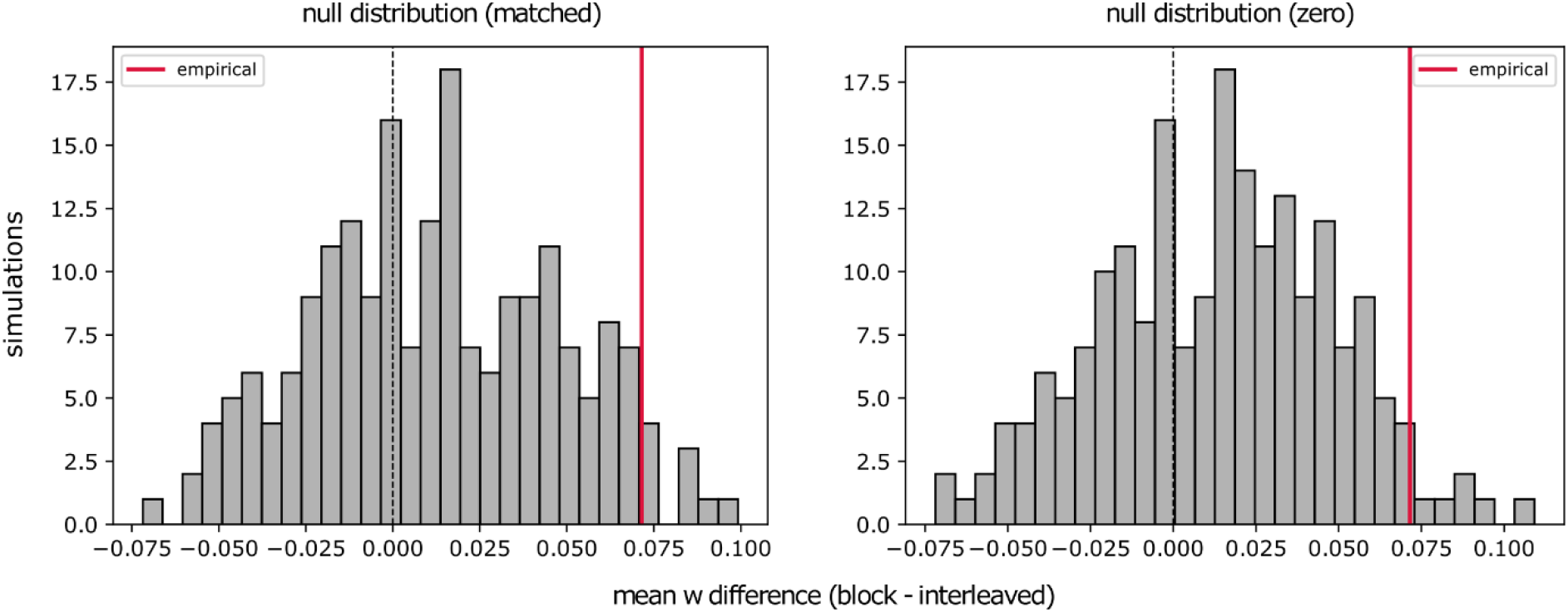
Parametric simulations controlling for potential bias. Histograms showing the empirical differences in w across conditions (red line, showing the paired statistics for participants with both block and interleaved trials) against the two null distributions generated from a participant-matched, across-condition mean w simulation (left) and a w=0 simulation (right).

**Figure S6.**
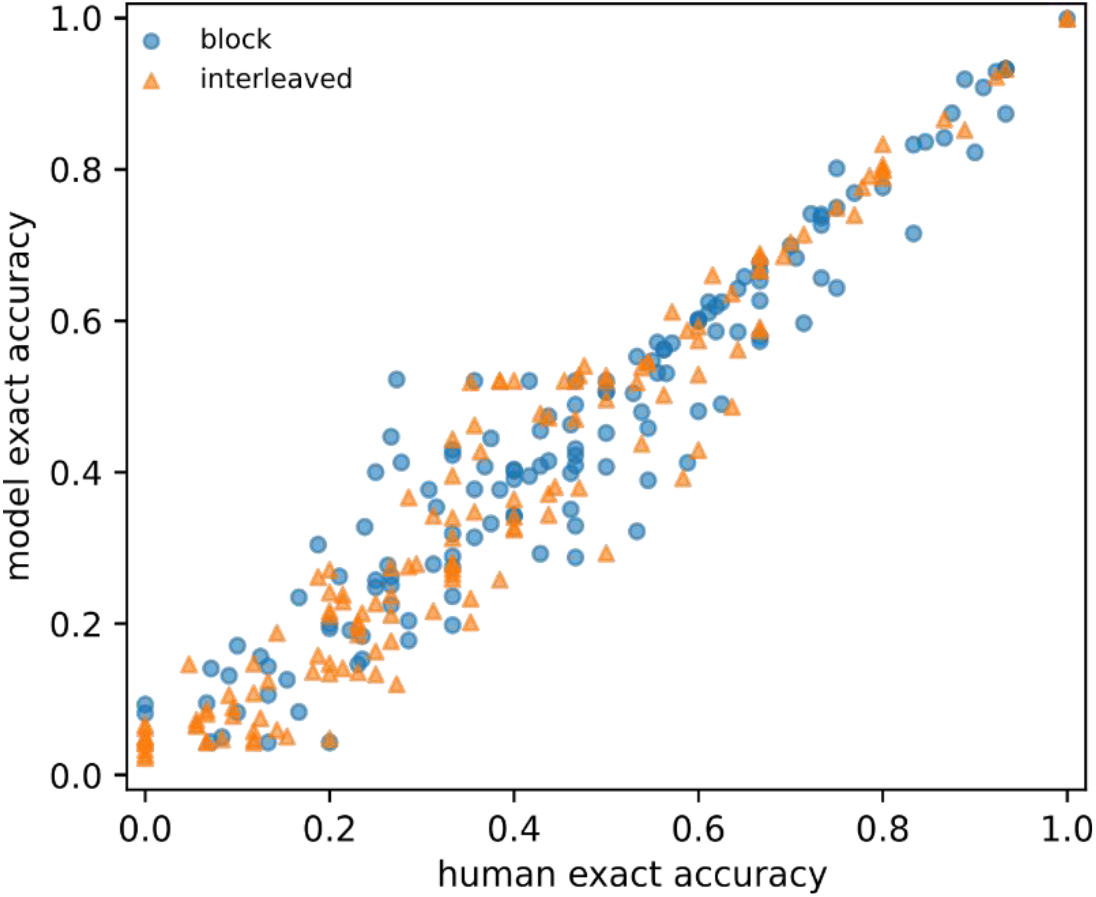
Model-simulated accuracy tracks individual differences in behavioural accuracy. Scatter plot shows empirical human accuracy versus accuracy computed under the fitted choice rule. We compared per-participant exact-ranking accuracy from the forward simulation against observed behavioural accuracy, for the block and interleaved conditions. Each point is one participant.

**Figure S7.**
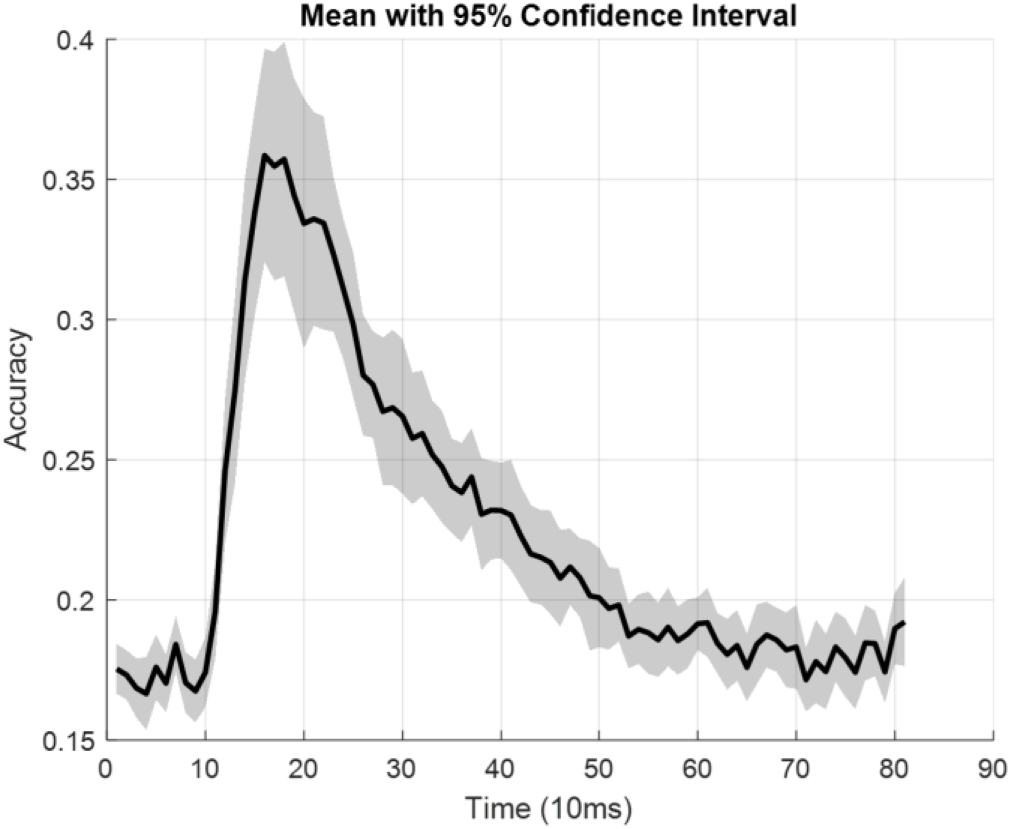
Cross-validation of stimulus decoder. Line chart showing the decoding accuracy of stimulus identity (chance level: 1/6 or 16.67%) when the stimulus (image) was shown on the screen (x-axis represents time after image cue onset). Shaded area represents 95% confidence interval and the black line represents the mean accuracy across the 25 participants.

**Figure S8.**
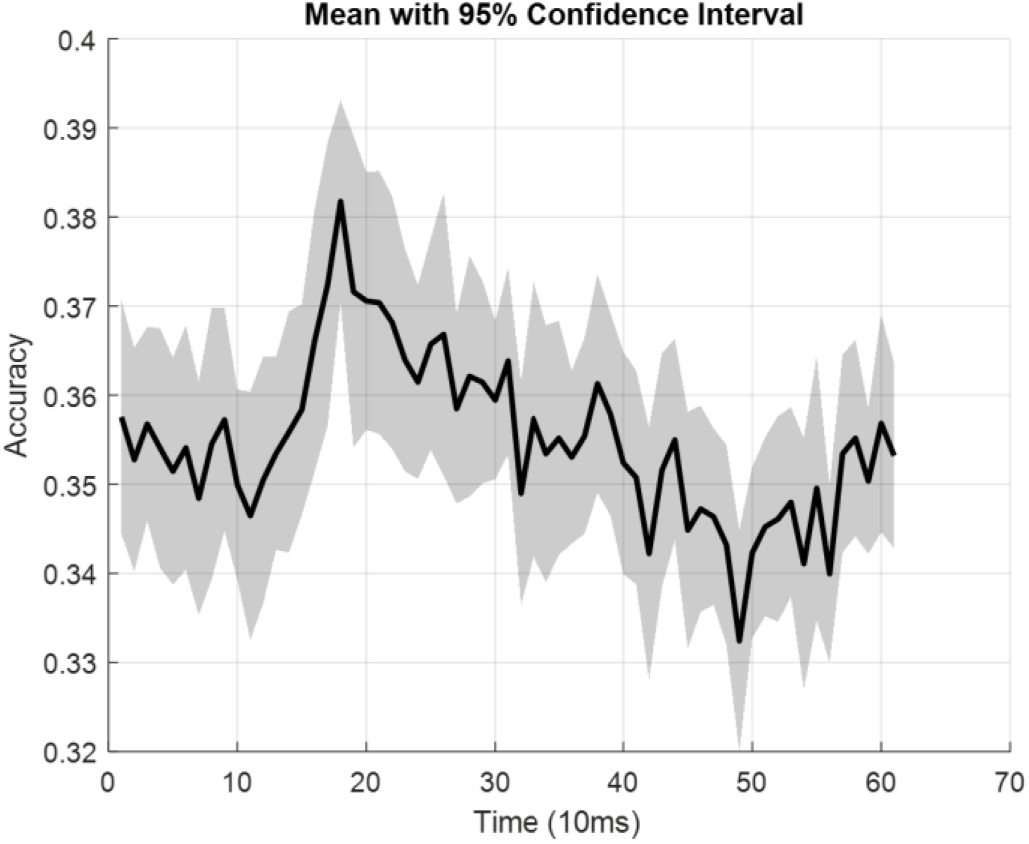
Cross-validation of subtask decoder. Line chart showing the decoding accuracy of subtask identity (chance level: 1/3 or 33.33%, with an above-chance baseline expected as the participant would know what subtask is coming next) when the word (text) was shown on the screen (x-axis represents time after word cue onset). Shaded area represents 95% confidence interval, and the black line represents the mean accuracy across the 25 participants.

**Figure S9.**
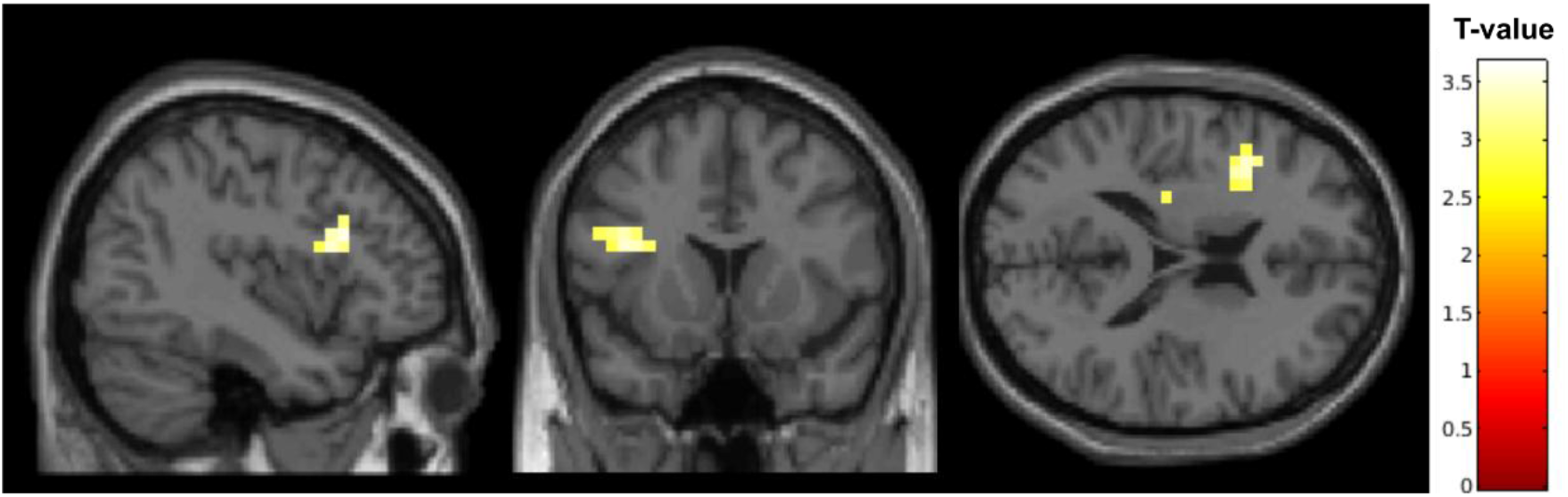
Source localisation revealed greater inferior frontal gyrus (IFG) source power during block planning. In an exploratory analysis, we examined source-localised broadband signal power during planning to ask whether a candidate brain region had greater involvement for block compared to interleaved condition. We focused on left IFG as our a priori region of interest, given its involvement in both hierarchical planning and structured task control (Jeon and Friederici, 2013; Koechlin and Jubault, 2006; Thompson-Schill et al., 1997). We hypothesised greater IFG source-level activity for the block compared to interleaved condition. Within an anatomically defined left IFG ROI (AAL3 atlas), we applied threshold-free cluster enhancement (TFCE; a multiple-comparisons correction method) with 5,000 permutations (including small-volume correction). In keeping with our hypothesis, a significant IFG cluster with enhanced broadband power (1-45 Hz, frequency range following Wimmer et al., 2020) was evident for block compared to interleaved planning (peak t= 3.51, FWE-corrected p = 0.027; peak MNI: −40, 19, 23). The brain map shows a whole-brain t-statistic (block > interleaved) from the LCMV beamformer analysis, thresholded at p < 0.005 uncorrected. Colour scale indicates t-values.

**Figure S10.**
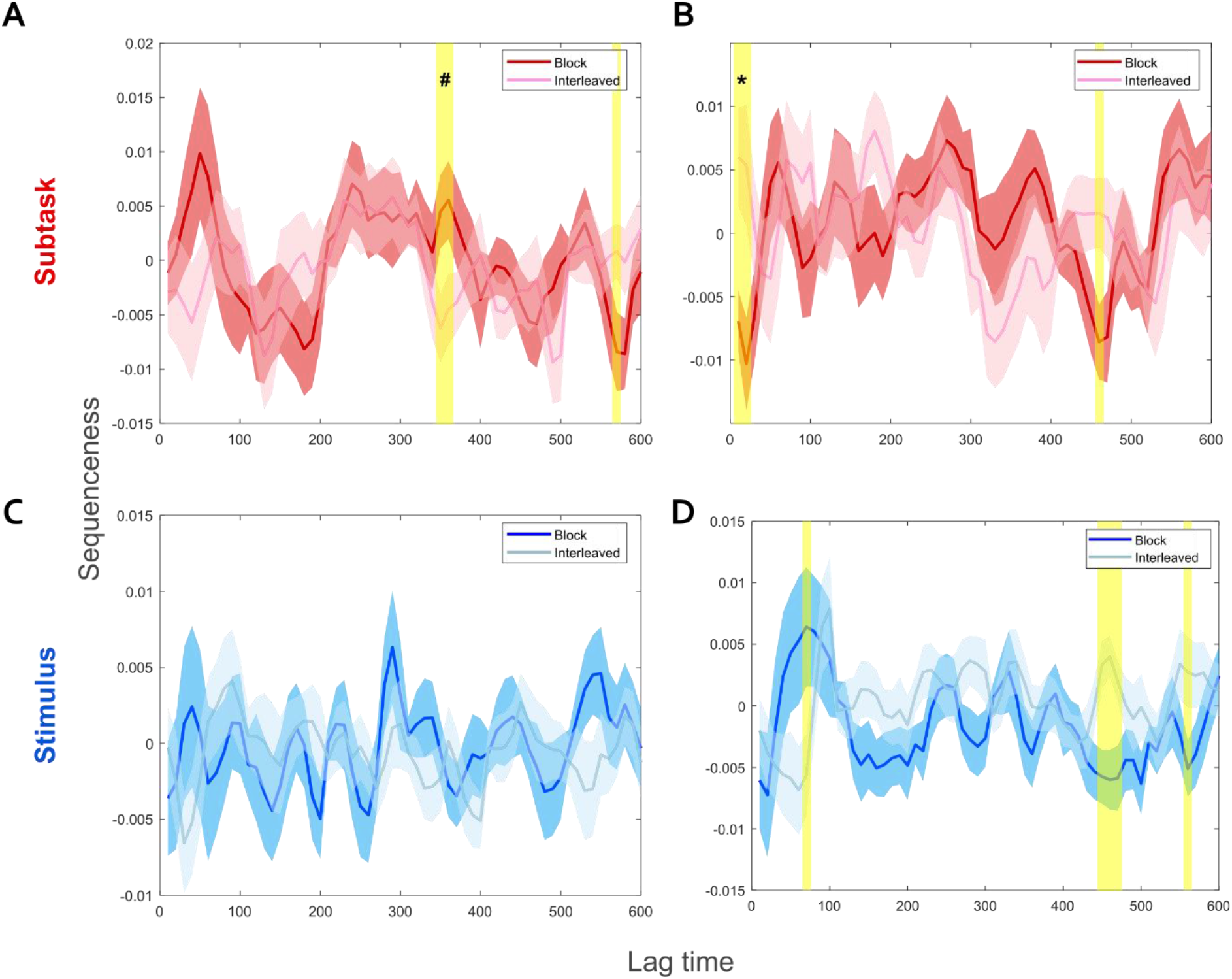
Differential stimulus and subtask replay in the block and interleaved conditions. (A) For the subtask forward replay, there is marginally higher sequenceness in the block condition than in the interleaved condition (p=0.0779, with 1,000 permutations of block/interleaved labels, FWE-corrected p-value for finding significant cluster mass thresholding at p=0.05 two-tailed t-stats). (B) On the other hand, for the subtask backward replay, there is significantly higher sequenceness in the interleaved condition than in the block condition (p=0.0370, with 1,000 permutations of block/interleaved labels, FWE-corrected p-value for finding significant cluster mass thresholding at p=0.05 two-tailed t-stats). (C) Consistent with the analyses in Fig. 5A that no significant forward stimulus replay was identified, we also did not find any significant differences between block and interleaved conditions for forward stimulus replay sequenceness (with no time point passing the p=0.05 t-statistics threshold). (D) For the stimulus backward replay, there are no significant differences in sequenceness between the interleaved and block conditions (p=0.1119, with 1,000 permutations of block/interleaved labels, FWE-corrected p-value for finding significant cluster mass thresholding at p=0.05 two-tailed t-stats). The shaded area surrounding the line represents the 95% confidence interval.

**Figure S11.**
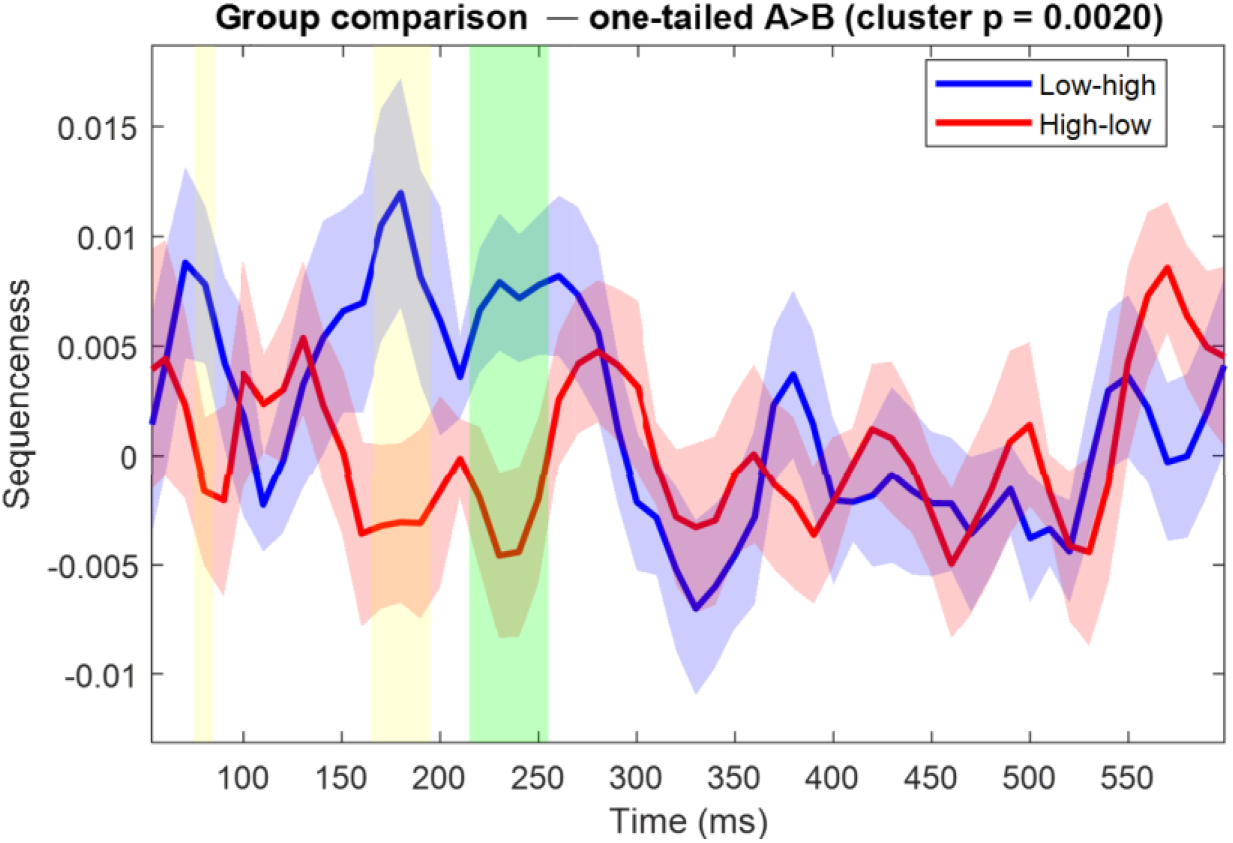
Replay with different ranking rules. For the subtask backward replay, there is significantly higher sequenceness in the low-to-high than in the high-to-low situation. Cluster-based permutation was performed with 1,000 permutations of labels, and the FWE-corrected p-value was used to find significant cluster mass thresholding at p=0.05 using one-tailed t-statistics. Shaded areas represent the clusters passing the threshold, and the one in green represents the strongest one (one-tailed p=0.002 in the cluster-based permutation test).

## Supplementary text

### Testing reliability of model comparison tools using model recovery

We considered six candidate models. The mixture model combines stimulus-level and subtask-level value with a free weight w. Two nested restrictions ask whether the arbitration between the two value sources is needed at all: w = 0, in which choice is governed only by stimulus-level value and corresponds to the literature standard, a classic Q-learner; and w = 1, in which it is governed only by subtask-level value. Each of these three models was also fitted with an additional lapse parameter, so that a proportion ε of rankings is drawn uniformly rather than from the learned values, giving P(ranking) = (1 - ε) * P (ranking | value) + ε / 4!. In these supplementary analyses, all models were fitted per participant by maximum likelihood, with the two conditions fitted jointly unless stated otherwise.

Before comparing models, we asked which selection criterion is capable of distinguishing them under this design. For each of the 238 participants, we simulated 100 datasets from each candidate model, using that participant’s trial sequence and maximum-likelihood parameters, refitted every candidate to every synthetic dataset, and asked how often each criterion identified the generating model. Each criterion needs a rule for deciding which model generated a given simulated dataset. For the Akaike Information Criterion corrected (AICc) and Bayesian Information Criterion (BIC), the rule is simply to select the model with the lower score. For the parametric bootstrap cross-fitting method (PBCM; Wagenmakers et al., 2004), the rule is based on d = LL(reference) - LL(rival), the difference in log-likelihood between the two fitted models: simulating from each model in turn yields two distributions of d, and the threshold is the value of d that best separates them, taken as the value maximising the sum of sensitivity and specificity. For the bootstrap likelihood ratio test (BLRT; Nylund et al., 2007), the rule is a critical value on the likelihood ratio statistic, taken as the 95th percentile of its null distribution simulated from the restricted model. This threshold and this critical value were estimated on one half of the simulated datasets and applied to the other, so that no criterion was evaluated on the data used to calibrate it.

Balanced accuracy (pooled across participants) favoured PBCM for every comparison (Table S1). BLRT was intermediate, and AICc and BIC were poorest throughout. The same ordering held for the lapse variants.

**Table S1.**
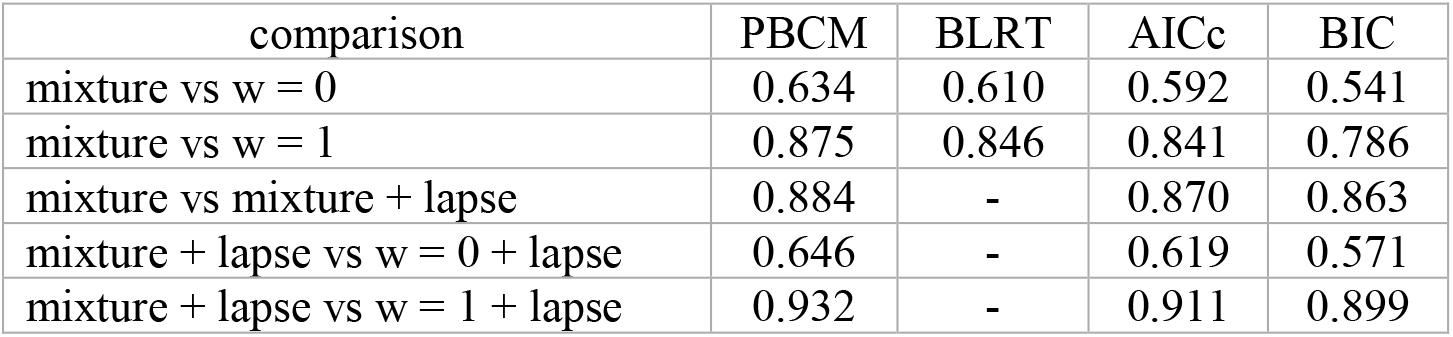
Balanced accuracy of each selection criterion at identifying the true data-generating model for simulated task behaviour, pooled across participants. BLRT is defined only for nested pairs within the original model set and was not computed for the lapse comparisons.

| comparison | PBCM | BLRT | AICc | BIC |
| --- | --- | --- | --- | --- |
| mixture vs $w = 0$ | 0.634 | 0.610 | 0.592 | 0.541 |
| mixture vs $w = 1$ | 0.875 | 0.846 | 0.841 | 0.786 |
| mixture vs mixture + lapse | 0.884 | - | 0.870 | 0.863 |
| mixture + lapse vs $w = 0$ + lapse | 0.646 | - | 0.619 | 0.571 |
| mixture + lapse vs $w = 1$ + lapse | 0.932 | - | 0.911 | 0.899 |

The differences in balanced accuracy understate how poorly AICc and BIC perform in model recovery, because the errors were strongly asymmetric (Table S2). For the comparison between the mixture model and the w = 0 restriction, AICc and BIC identified the w = 0 model almost perfectly when it had generated the data (specificity 96.8% and 99.6%, respectively) but identified the mixture model in only 21.6% (AICc) and 8.6% (BIC) of cases when the mixture had generated it. Cross-fitting every candidate model to data generated by the candidate model, PBCM was approximately unbiased (sensitivity 63.2%, specificity 63.6%). The same bias towards the simpler model appeared in every comparison involving a lapse: when the four-parameter (without lapse) and five-parameter mixture (with lapse) models were compared, AICc and BIC identified the four-parameter model on 94.5% and 98.3% of occasions when it generated the data but identified the five-parameter model on only 79.5% and 74.3%, whereas PBCM was again close to symmetric and correctly identified the four-parameter and five-parameter models in 92.1% and 84.6% of cases, respectively. We therefore base the model comparison on the bootstrap procedures.

**Table S2.** Sensitivity and specificity of each criterion. Sensitivity is the probability of identifying the reference model when it generated the data; specificity is the probability of identifying the alternative.

| comparison |  | PBCM | BLRT | AICc | BIC |
| --- | --- | --- | --- | --- | --- |
| mixture vs w = 0 | sensitivity | 0.632 | 0.286 | 0.216 | 0.086 |
|  | specificity | 0.636 | 0.934 | 0.968 | 0.996 |
| mixture vs w = 1 | sensitivity | 0.887 | 0.759 | 0.707 | 0.574 |
|  | specificity | 0.864 | 0.932 | 0.974 | 0.998 |
| mixture vs mixture + lapse | sensitivity | 0.921 | - | 0.945 | 0.983 |
|  | specificity | 0.846 | - | 0.795 | 0.743 |
| mixture + lapse vs w = 0 + lapse | sensitivity | 0.640 | - | 0.286 | 0.149 |
|  | specificity | 0.651 | - | 0.953 | 0.993 |
| mixture + lapse vs w = 1 + lapse | sensitivity | 0.937 | - | 0.860 | 0.803 |
|  | specificity | 0.927 | - | 0.962 | 0.994 |

### Comparison of the mixture model against alternatives

We compared the mixture model against the nested restrictions w = 0, in which choice is governed only by stimulus-level value, and w = 1, in which it is governed only by subtask-level value.

Subtask-only learning (w = 1) was rejected. This model cannot account for the ordering of stimuli within a subtask, because it assigns every stimulus in a subtask the same value. Cross-fitting assigned 230 of 238 participants to the mixture model (Table S3; the log-likelihood difference summed across participants, 2499.9, against a criterion of 1289.4), and the bootstrap likelihood ratio test rejected the restriction decisively (Λ, the likelihood ratio statistic 2[LL(mixture) - LL(restricted)] summed across participants = 4999.7, against a 95% critical value of 177.0 obtained from the simulated null, p = 0.0099, the smallest value attainable with 100 bootstrap samples; median per-participant Λ = 11.18 against a null median of 0.02; 179 of 238 participants individually significant at p < 0.05).

Stimulus-only learning (w=0) was also rejected. This model does not have the ability to account for any influence of subtask structure on choice, because it learns each stimulus independently. Cross-fitting assigned 152 of 238 participants to the mixture model against 86 to the restriction, and the group-level statistic exceeded its bootstrap criterion (220.3 against 194.5). The bootstrap likelihood ratio test rejected the restriction (Λ summed across participants = 440.6, against a 95% critical value of 205.5, p = 0.0099; median per-participant statistic 0.49 against a null median of 0.04; 36 of 238 participants individually significant, against the 12 expected by chance at p < 0.05).

### Comparison of the mixture model with and without a lapse parameter

We next compared the mixture model with and without a lapse parameter (four vs five parameters). Cross-fitting assigned 215 of 238 participants to the five-parameter model against 23 to the four-parameter model (Table S3). Correcting the observed proportions for the bootstrap error rates suggested that all estimated participants favouring the five-parameter model, with the group-level statistic also favoured the five-parameter model. The bootstrap likelihood ratio test rejected the four-parameter model (Λ = 4595.6 against a 95% critical value of 107.0, p = 0.0099; median per-participant Λ = 15.68 against a null median of 0.00; 194 of 238 participants individually significant). The same conclusion was reached separately in each condition. Because AICc and BIC are biased towards the simpler model in this design (Table S2), the fact that the richer model wins under the bootstrap procedure is evidence against the direction of that bias rather than with it; here the three criteria nonetheless agreed on 90.9% of participants for AICc and 84.0% for BIC.

### Comparison within the five-parameter model set

We repeated the comparison against the two restrictions with a lapse parameter added to each, fitting the two conditions jointly (n = 238; Table S3). Subtask-only learning was again rejected decisively: cross-fitting assigned 226 of 238 participants to the mixture model, the corrected proportion favouring it was 1.000 (95% CI [0.98, 1.00]), and the bootstrap likelihood ratio test rejected the restriction (Λ = 8729.4 against a 95% critical value of 224.5, p = 0.0099; median per-participant Λ = 28.45 against a null median of 0.03; 205 of 238 participants individually significant).

Stimulus-only learning remained the harder comparison. Cross-fitting assigned 148 of 238 participants to the mixture model against 90 to the restriction, and the group-level statistic exceeded its bootstrap criterion only narrowly (255.4 against 251.8). The corrected proportion favouring the mixture model was 0.836 (95% CI [0.66, 1.00]). The bootstrap likelihood ratio test rejected the restriction (Λ = 510.8 against a 95% critical value of 201.6, p = 0.0099; median per-participant statistic 0.40 against a null median of 0.00; 45 of 238 participants individually significant). This mirrors the four-parameter comparison, in which the same restriction gave the narrowest margin, and reflects the limited power of the design to separate a small mixture weight from none at all.

**Table S3.** Model comparison on the observed data. Cross-fitting assignments, the prevalence-corrected proportion of participants favouring the richer model.

| comparison | assigned to the richer model | corrected proportion [95% CI] | likelihood ratio statistic $\Lambda$ (critical value) |
| --- | --- | --- | --- |
| mixture vs $w = 1$ | 230/238 | 1.000 [1.00, 1.00] | 4999.7 (177.0) |
| mixture vs $w = 0$ | 152/238 | 0.881 [0.70, 1.00] | 440.6 (205.5) |
| mixture + lapse vs mixture | 215/238 | 1.000 [1.00, 1.00] | 4595.6 (107.0) |
| mixture + lapse vs $w = 1$ + lapse | 226/238 | 1.000 [0.98, 1.00] | 8729.4 (224.5) |
| mixture + lapse vs $w = 0$ + lapse | 148/238 | 0.836 [0.66, 1.00] | 510.8 (201.6) |

### Model selected

The mixture model with a lapse parameter was selected over the four-parameter mixture model and over both restrictions within the five-parameter set. The two restrictions were rejected under both the four-parameter and the five-parameter model, so the conclusion that choice draws on both stimulus-level and subtask-level value does not depend on which is used. We therefore used the five-parameter (fitting both lapse and w) model in all other analyses in the paper.

